# UV-B-Induced DNA Repair Mechanisms and Their Effects on Mutagenesis and Culturability in *Escherichia coli*

**DOI:** 10.1101/2024.11.14.623584

**Authors:** Sreyashi Ghosh, Jenet Narzary, Mehmet A. Orman

**Author notes:** Correspondence to: S222 Engineering Bldg 1, 4226 Martin Luther King Boulevard, Houston, TX 77204-4004.

## Abstract

Ultraviolet-B (UV-B) radiation, intensified by ozone depletion, induces DNA damage and promotes mutagenesis, shaping evolution. While UV-induced SOS responses are well characterized in bacteria, the cellular consequences of prolonged UV-B exposure remain less clear. Prolonged UV-B exposure disrupts translation and RecA-mediated SOS induction without major changes in membrane permeability or reactive oxygen species. This impairs mutagenesis and induces a reversible loss of culturability. Genetic analysis reveals both redundant and differential roles for DNA repair pathways: homologous recombination (RecA, RecB), nucleotide excision repair (UvrA), and translesion synthesis (UmuC/D) are essential for maintaining mutagenesis and culturability, while others (RecN, RmuC) have limited impact. Notably, deletion of UvrD (a repair-associated helicase) intensifies transient non-culturability without affecting mutagenesis, underscoring complexity in repair networks. Overall, our findings reveal a dose-dependent trade-off: moderate UV-B promotes mutagenesis with minimal viability loss, whereas prolonged exposure suppresses mutagenesis via transient dormancy, reflecting an adaptive strategy with significant evolutionary implications.

## Introduction

Mutagenic processes are key to evolutionary progress ^1–3^, as they generate genetic diversity. This is essential for organisms to adapt to and endure in dynamic settings that are continuously changing. Understanding the mechanisms underlying mutagenesis helps us better comprehend the evolutionary forces shaping organisms ^4,5^. This knowledge also aids in developing more potent and effective medical treatments to combat many diseases, given that pathogenic microorganisms and tumorigenic cells develop resistance through mutagenesis^6–9^.

Ultraviolet (UV) radiation, particularly UV-B, plays a dual role in Earth’s ecosystems; it acts as both a developmental cue and an environmental stressor. While moderate UV-B levels may regulate processes like plant defense and microbial interactions, elevated UV-B due to ozone depletion has been linked to ecological disruptions and even mass extinction events, such as the end-Permian and Devonian–Carboniferous extinctions^10–13^. UV-B is of particular interest because it is thought to reach the Earth’s surface in greater intensity during periods of severe ozone depletion caused by catastrophic events. On a microbial scale, UV radiation has continually shaped the trajectory of evolution over billions of years, from contributing to the origins of life to driving genetic variation and adaptation^14,15^. This is because UV radiation directly damages DNA and induces certain repair mechanisms that are highly mutagenic^16–19^. The UV light spectrum, ranging from 100 to 400 nanometers (nm), can be categorized into four groups: UV-A (315-400 nm), UV-B (280-315 nm), UV-C (100-280 nm), and vacuum-UV (100-200 nm)^20^. UV-C is more readily absorbed by nucleic acids compared to UV-B and UV-A^21^. When bacteria are exposed to UV radiation, ^6,7^ it triggers the formation of pyrimidine dimers and other DNA lesions that disrupt the normal structure and function of DNA molecules^22,23^. These UV lesions are removed by various effective bacterial systems, such as the photoreactivation repair system (i.e. dimer monomerization under visible light by photolyase enzymes, such as Phr)^24^, base excision repair (BER) (removal of cyclobutane-thymine dimers using DNA glycosylases and AP endonucleases)^25^, and the primary pathway for removal of bulky DNA lesions, i.e. the nucleotide excision repair (NER) (for rapid repair of photo-lesions using the *uvr* genes, *polA* and ligase enzyme) ^26^. However, an excessive amount of dimers that cannot be removed by the repair systems can accumulate and interfere with the cellular replication process, resulting in the formation of “secondary” lesions in the form of ssDNA fragments^27^. Therefore, to deal with this severe DNA damage, a post-replication damage repair mechanism known as the bacterial SOS response pathway is induced, preventing premature cell division ^16,28,29^ and providing cells with sufficient time to repair the damaged DNA^18,30–32^. This SOS response in bacteria is controlled by the multifunctional RecA protein, a key component that cleaves the transcriptional repressor LexA^33–35^. This cleavage initiates the expression of more than 40 SOS genes. RecA, involved in the recombination repair of ssDNA gaps^36^, helps provide an error-free damage removal route. However, excessive DNA damage leads to the induction of the mutagenic phase of the SOS response, mediated by bacterial DNA polymerase enzymes, including Pol V ^37,38^, which is encoded by the *umuD* and *umuC* genes ^39^. Pol V can bypass template lesions during DNA replication through a process known as translesion DNA synthesis (TLS) ^40^. Two additional TLS DNA polymerases, Pol II (encoded by *polB*) and Pol IV (encoded by *dinB*), facilitate replication past blocking lesions, albeit at the cost of introducing mutations. Another DNA damage repair strategy is the mismatch repair (MMR) pathway ^41^, which recognizes base-base mismatches and insertion/deletion loops, including those introduced by DNA polymerases. However, with increased mutagenic stress, the efficiency of MMR decreases, as observed in the case of fluoroquinolone treatment ^42^. Such stress can potentially activate the SOS response and be addressed by SOS-mediated repair mechanisms, as the accumulation of DNA lesions and stalled replication forks are known to trigger this global DNA damage response.

UV radiation can serve as a valuable experimental tool for investigating mutagenesis because it induces a wide range of mutations without an apparent sequence preference^43,44^. Additionally, UV radiation is of great interest to scientists for its applications in biotechnology, particularly in directed evolutionary strategies to engineer proteins or organisms^45,46^. Moreover, there is growing interest in using UV radiation as a disinfectant, including for sterilizing air, equipment, and surfaces; reducing the transmission of airborne diseases; and even treating wound infections^47,48^. Despite their increased applications in clinical settings, we believe that two phenomena, UV-induced bacterial cell dormancy and mutagenesis, need to be fully understood. Bacterial cells are highly heterogeneous, and they can enter a growth-arrested state through stochastic mechanisms or environmental factors^49^, including UV treatment, which can make these cells highly tolerant. Also, the SOS-response-mediated cell dormancy not only promotes cell survival but also sustains mutagenic processes, as dormant cells may still harbor DNA damage^50,51^. Additionally, UV radiation itself is highly mutagenic; it can accelerate the emergence of more resilient mutants, which could pose a serious threat to public health.

UV-induced repair mechanisms are highly complex as UV can damage many cellular components, including DNA, RNA, lipids, and proteins^52,53^. While the role of the SOS response in mutagenesis has been extensively studied (refer to the following review articles^16,30,32^ for more details), our understanding of the downstream repair mechanisms and their contributions to cell survival, culturability, and mutagenesis remains limited. Although our recent study showed an almost perfect correlation between RecA levels and mutation frequency in *E. coli* following UV-B treatment, prolonged UV-B exposure was found to impair SOS-mediated mutagenesis and induce a transient non-culturability state ^54^. While this may represent a potential survival strategy for cells to evade UV-B-induced mutagenesis, the underlying mechanisms remain unclear. Our current study aims to investigate these complex mechanisms further, providing new insights into the molecular processes underlying mutagenesis and dormancy.

## Results and Discussion

### The transient unculturability is largely driven by the global regulator RecA, crucial for the SOS response and recovery

To study UV-mediated mutagenesis and the SOS response, we used our previously established methodology^54^. We cultured *E. coli* MG1655 wild-type cells in 2 mL Lysogeny Broth (LB) medium in test tubes until they reached the mid-exponential phase with an optical density (OD_600_) of ~0.5. Subsequently, the cells were transferred to petri dishes, forming a thin film of cultures that increased the surface area for UV exposure. This film was then subjected to UV-B light (302 nm thin-line transilluminator, UVP ChemStudio, Analytik Jena) for various durations: 2, 4, 8, 16, 24, and 32 minutes (min), ensuring a wide range of UV intensity from 120 J/m^2^ to as high as 1920 J/m^2^ as shown in **Fig 1a** (see **Supplementary Fig. S1,** and **Materials and Methods**). We used a broad UV-B dosage range to capture dose-dependent effects, as UV-B is less energetic than UV-C and requires higher exposure to induce cellular stress. This approach aligns with prior studies using UV-B, where treatment durations and doses vary widely depending on strain and experimental conditions^55–59^. Following exposure, the cells in LB were promptly transferred back to test tubes and cultured in a shaker for a 24-hour recovery (refer to the Materials and Methods section for details). Cultures that did not receive UV-B treatment served as controls. During recovery, colony-forming units (CFU) were quantified at the indicated time points for each condition, as shown in **Fig. 1b**. At the beginning of recovery (t~0), CFU levels decreased in cultures exposed to longer UV-B treatments. A 4-8 min exposure caused a 10-fold reduction, while 16 min led to a ~100-fold drop compared to the control. Exposure times of 24 and 32 min resulted in a ~10,000-fold reduction (**Fig. 1b**). After 15 minutes, CFU levels notably increased in cultures with 24 and 32 min of UV-B exposure, likely due to mechanisms making cells temporarily unculturable. This drastic increase in CFU levels is not due to cell division, given that the *E. coli* doubling time is about 20-25 minutes, which is too short for significant growth within that period. By the 24-hour mark, all conditions showed similar CFU levels (**Fig. 1b**).

**Fig. 1:**
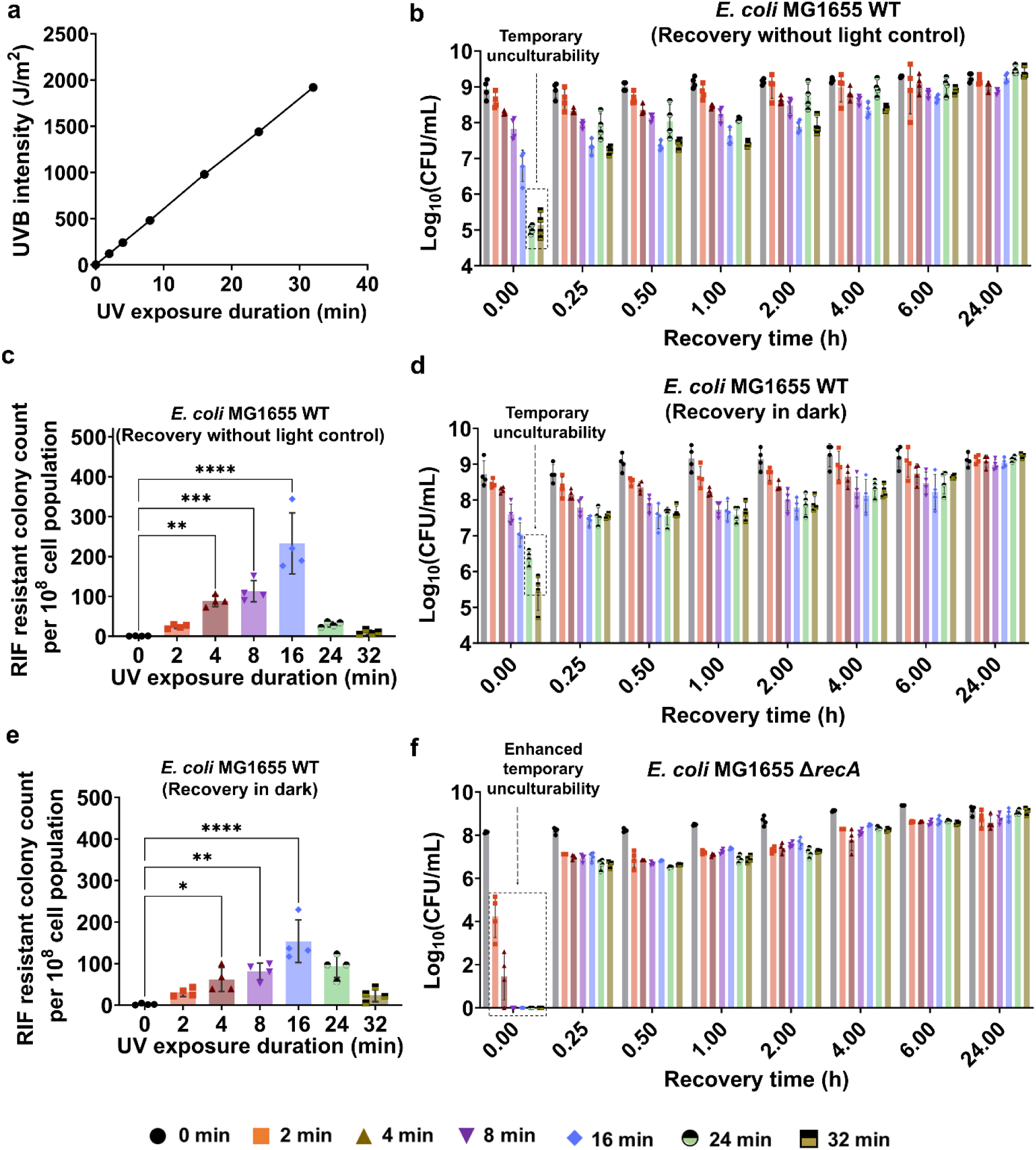
Excessive UV exposure resulted in transient loss of culturability and decreased mutagenesis. (a) The graph shows the relationship between UV-B exposure time (min) and calculated energy dosage (J/m^2^), based on measured irradiance (see Materials and Methods). (b) Exponential-phase *E. coli* MG1655 WT cells were exposed to UV-B light for 0, 2, 4, 8, 16, 24, and 32 minutes, followed by a 24-h recovery period. At specific time points during recovery (t = 0 h, 0.25 h, 0.5 h, 1 h, 2 h, 4 h, 6 h, and 24 h), cells were collected and plated to determine colony-forming units (CFU). (c) Levels of UV-induced rifampicin (RIF) resistance mutations were measured by counting RIF-resistant colonies (per 10^8^ cells) in the WT cultures after recovery for the indicated UV exposure times. (d) CFU counts and (e) mutation frequency of *E. coli* MG1655 following UV-B exposure were determined under dark conditions. The same experimental setup described above was used, but cultures were recovered in the absence of light to eliminate photoreactivation. (f) The temporal CFU profiles of *E. coli* MG1655 Δ*recA* cells were monitored during recovery following UV treatment. n=4. Statistical analysis was performed using one-way ANOVA with Dunnett’s post-test, where \**P* < 0.05, \*\**P* < 0.001, \*\*\**P* < 0.01, \*\*\*\**P* < 0.0001. Data corresponding to each time point represent mean value ± standard deviation.

To measure mutagenesis, we quantified rifampicin (RIF)-resistant colonies after 24 hours of recovery. Cultures were plated on LB agar with 500 µg/ml RIF, and colony counts were reported per 10^8^ cells (**Fig. 1c**). Although this method does not directly measure mutation levels, it is widely used ^44,60–62^ due to the strong correlation between mutagenesis and resistant colony formation ^63,64^. We chose a 24-hour recovery period to ensure accurate colony counts, consistent with previous studies^60,65,66^, as mutation levels generally plateau by this time^54^. Our results showed that mutagenesis increased with UV-B exposure, peaking at 16 minutes, but longer exposures (24-32 min) significantly reduced RIF-resistant colonies (**Fig. 1c**). This pattern, consistent with our previous findings^54^, suggests a link between UV-B exposure, mutagenesis, and temporary unculturability.

Our initial experiments were conducted without controlling for light exposure. Given the role of photoreactivation in bacterial recovery, we investigated whether the observed transient loss and restoration of culturability were driven by photolyase-mediated repair. The *phr* gene encodes DNA photolyase, which reverses UV-induced cyclobutane pyrimidine dimers in the presence of blue light^67,68^. As we were unable to obtain viable Δ*phr* knockouts in *E. coli* MG1655 using the λ-Red recombinase technique (see Materials and Methods)^69^, a result consistent with a previous report ^70^, we evaluated the role of photoreactivation by repeating the UV-B exposure and recovery experiments under strict dark conditions. Importantly, despite minor fluctuations in mutation frequency, expected due to altered photolyase-mediated repair mechanisms, we observed the same core features under dark conditions: a pronounced but transient decline in culturability, followed by recovery, and a dose-dependent mutation frequency (**Fig. 1d, e**). To directly test whether *phr* function is necessary for these outcomes, we conducted parallel experiments using a Δ*phr* strain from the Keio collection (*E. coli* BW25113 Δ*phr*), as this strain can have a viable knockout *phr* strain. Upon UV-B treatment and recovery, this strain exhibited a comparable transient unculturability and dose-dependent mutation frequency (**Supplementary Fig. S2**), mirroring the responses seen in MG1655. While some quantitative differences were observed in BW25113 (likely due to the distinct genetic background), these results indicate that the observed phenomena were not solely contingent on light-driven DNA repair. This is plausible, as UV-induced mutations likely arise from error-prone repair mechanisms rather than error-free photoreactivation by Phr.

In our previous study, we found that excessive UV-B treatment (24 and 32 minutes) impaired the SOS response, including the expression level of the RecA protein^54^. Therefore, the observed transient unculturability is likely more complex and may involve various mechanisms, given that the SOS response is intricate, involving multiple proteins, with RecA serving as the global regulator. We hypothesized that if this transient unculturability is linked to the impaired SOS response, then genetic perturbation of this response through *recA* deletion would further decrease cell culturability. Our results verified that the knockout strain of *recA* exhibited a drastic reduction in culturability immediately after UV treatment across all conditions (**Fig. 1f**). Initially, CFU levels in the Δ*recA* strain were below the limit of detection (1 CFU) for most UV treatment conditions; however, this was transient, as we observed a rapid increase in CFU levels (from 1 to 10^7^) within 15 minutes of recovery (**Fig. 1f**). This observation is highly interesting, and to our knowledge, such a dramatic recovery pattern in a *recA*-deficient background has not been previously reported. Although liquid-holding recovery has been well documented since the 1960s (primarily involving excision repair and photoreactivation during non-nutritive incubation in wild-type strains)^71,72^, our findings are distinct, given that the marked, transient increase in culturability following UV-B treatment occurs in a genetic background lacking a functional SOS response. While we previously demonstrated that prolonged UV-B exposure impairs the RecA-mediated SOS response and that *recA* deletion completely eliminates mutagenesis across all exposure durations (and thus did not repeat those experiments here)^54^, it remains unclear how downstream processes governed by RecA influence cell culturability and mutagenesis. We therefore aim to investigate these aspects in the following sections of this study.

### Genes significantly expressed during the SOS response have a moderate or negligible effect on cell culturability and mutagenesis

To identify the downstream mechanisms, we screened a subset of the *E. coli* promoter library containing 1900 promoter reporters. In this library, the promoters of *E. coli* genes are fused to a fast-folding green fluorescent protein (GFP) gene in low-copy-number plasmids, allowing accurate and reproducible measurement of gene expression^73^. We focused on approximately 40 reporters containing promoters that regulate the key SOS response and associated genes. These promoters are under the control of the RecA-LexA regulatory system, chosen based on a comprehensive analysis of *E. coli* databases^74^. In our experiments, *E. coli* MG1655 cells containing the selected promoters were cultured individually until the mid-exponential phase in 96-well plates. Subsequently, they were exposed to UV-B radiation for 16 min. This duration was chosen as it induces maximum SOS upregulation and mutagenesis without drastically impacting cell culturability (**Fig. 1b,c**). Also, 16-min UV treatments were used as positive controls for the subsequent sections. GFP levels were measured after 24 h of recovery, allowing adequate time for UV-induced promoters to express GFP. High cell densities ensured reliable GFP measurements, given that we used a plate reader in this screening assay.

Our specific focus was on genes showing significant upregulation following UV treatment (**Fig. 2a**), as they likely play pivotal roles in the downstream mechanisms of RecA-mediated SOS response. Several promoters showed drastic upregulation, including P*_recA_* and P*_lexA_* (promoters of genes encoding SOS response global regulators, as expected), P*_recN_* and P*_rmuC_* (promoters of genes involved in DNA recombinational repair), P*_sulA_* (promoter of a gene encoding cell division inhibitor), P*_polB_* and P*_dinB_* (promoters of genes encoding DNA polymerase enzymes), P*_ftsK_* (promoter of a gene encoding essential cell division protein), P*_sbmC_* (promoters of a gene encoding inhibitor of DNA gyrase-mediated DNA supercoiling), and P*_ybfE_* (promoter of a *lexA-*regulated gene whose function is not well characterized)^75^ (**Fig. 2a**). To determine whether the genes associated with the identified promoters are involved in SOS response-mediated culturability and mutagenesis, we generated single deletions in *E. coli* MG1655 for these genes (except for *lexA* and *ftsK*, as they are essential and could not be deleted^76–78^). The knockout strains were exposed to UV-B for 16 min or 32 min or untreated, and their CFU profiles during recovery, as well as their RIF-resistant colony levels, were similarly measured. For all knockout strains, the temporal CFU profiles remained the same as that of the wild-type, showing a transient non-culturability during the first 15 min of recovery after a high UV-B exposure duration of 32 min (**Fig. 2b, Supplementary Fig. S3**). As expected, the deletion of *recA* completely eliminated mutagenesis (**Fig. 2c**). Although 32 min of UV-B exposure significantly reduced mutagenesis in all strains, including wild type, the two DNA recombination gene knockouts, Δ*recN* and Δ*rmuC*, showed a 3-fold and 1.5-fold decrease in RIF-resistant mutant levels, respectively, compared to WT cells following 16 min of UV-B treatment (**Fig. 2c**). Surprisingly, other single deletion strains, including those for the genes encoding DNA polymerase II and IV, *dinB* and *polB*, did not show significant changes (**Fig. 2c**).

**Fig. 2:**
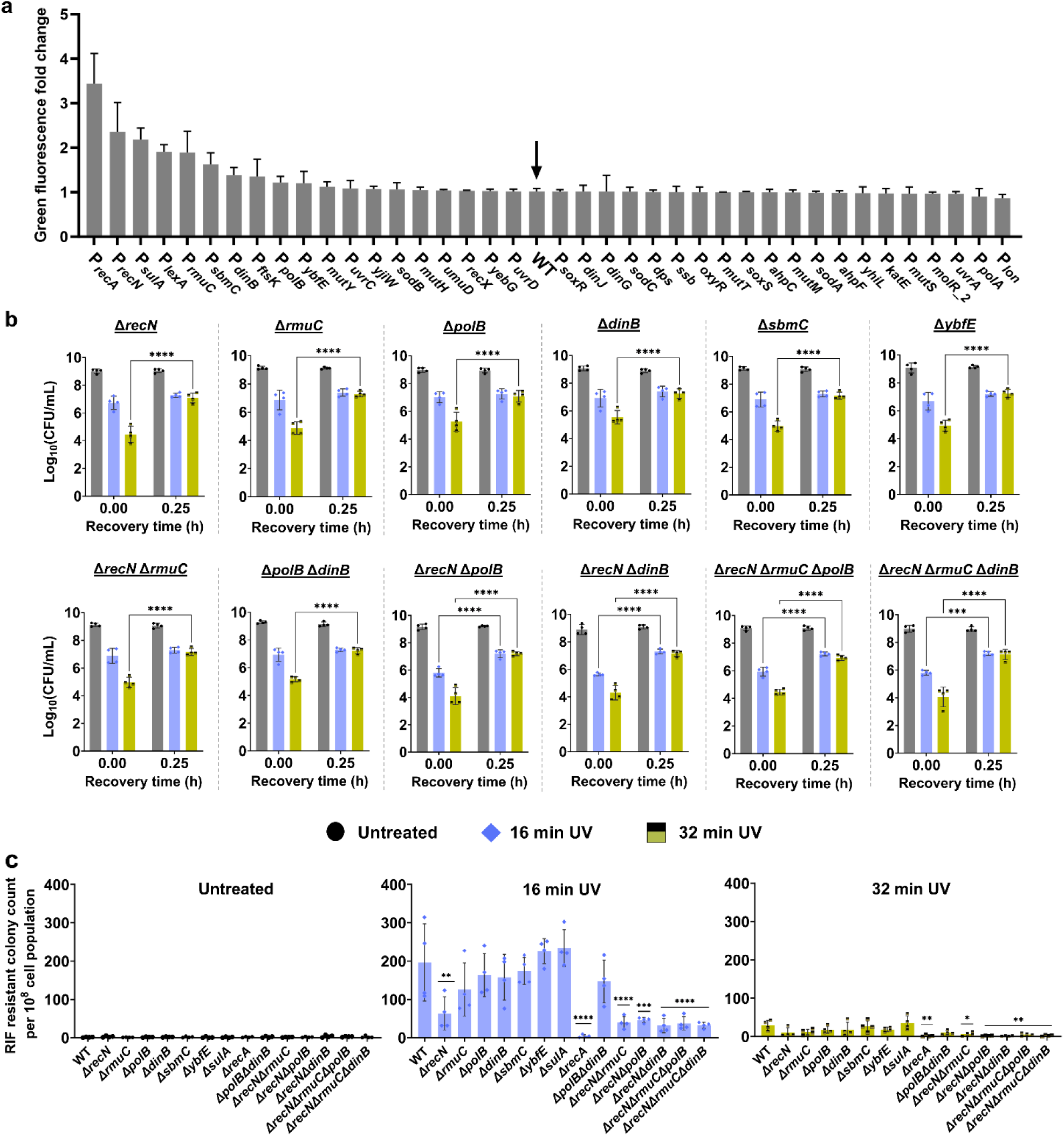
UV-induced upregulation of SOS genes and its impact on mutagenesis and cell culturability. (a) A library of *E. coli* MG1655 strains with promoter reporters for SOS response genes was UV-treated during the mid-exponential phase and allowed to recover for 24 h. GFP levels were measured after recovery and normalized to untreated controls. WT cells without any promoter reporters were used as background controls. (b) Genes upregulated following UV exposure were individually or combinatorially deleted in *E. coli* MG1655, and a mutagenesis assay was performed. Mid-exponential phase cells were exposed to UV for 0, 16, and 32 min and recovered for 24 h. At specified time points during recovery (t = 0 h, 0.25 h, 0.5 h, 1 h, 2 h, 4 h, 6 h, and 24 h), cells were collected and plated to determine CFU. Note that only the 0 and 0.25 h time points are shown here; the full-time course is provided in **Supplementary Fig. S3**. (c) RIF-resistant cells were quantified by plating samples on RIF-agar plates after 24 h of recovery, with results reported as RIF-resistant colony counts per 10^8^ cells for different UV exposure durations. n=4. Statistical analysis was performed using one-way ANOVA with Dunnett’s post-test, where \**P* < 0.05, \*\**P* < 0.001, \*\*\**P* < 0.01, \*\*\*\**P* < 0.0001. Data corresponding to each time point represents mean value ± standard deviation.

Overall, our results showed that single deletions of the downstream genes of RecA did not drastically impact cellular culturability and mutagenesis, except for RecN and RmuC deletions, which slightly reduced mutagenesis. RecN is involved in the recombinational repair of DNA double-strand breaks, and its mutation makes cells sensitive to mitomycin C and ionizing radiation^79–81^. Although RmuC is thought to be an inner membrane protein with a nuclease domain^82,83^, its function is not well understood, and our data indicate it only moderately impacts UV-B-mediated mutagenesis. It is quite surprising that the deletion of DNA polymerase II and IV, two key proteins involved in indirect mutagenesis^84^, did not impact UV-B-induced mutagenesis. As mutagenesis is a multifaceted process regulated by multiple mechanisms, testing multi-deletion strains might be necessary to show the complex interplay between these mechanisms. Furthermore, deleting a single polymerase enzyme may have minimal impact due to the abundance of the other polymerase enzyme. Therefore, we first generated Δ*recN*Δ*rmuC* and showed that the double deletion resulted in a cumulative decrease (~5 fold) in RIF-resistant colony levels compared to the wild type following 16 min of UV-B treatment (**Fig. 2c**). To assess the impact of DNA polymerase II and IV, we generated a Δ*polB*Δ*dinB* mutant strain, which did not affect RIF-resistant colony levels (**Fig. 2c**). Additionally, we generated Δ*recN*Δ*polB* and Δ*recN*Δ*dinB* double knockout strains and demonstrated that they exhibited approximately a 5-fold reduction in RIF-resistant colony levels compared to wild-type cells (**Fig. 2c**). Deletion of DNA polymerase genes *polB* and *dinB* from Δ*recN*Δ*rmuC* individually did not show significant changes compared to the Δ*recN*Δ*rmuC* strain in RIF-resistant mutant levels (**Fig. 2c**), suggesting that the observed reduction in RIF-resistant colonies in multi-deletion strains is primarily due to the effects of *recN* and/or *rmuC*.

### Screening the knockout strains reveals the redundancy of repair mechanisms in UV-induced mutagenesis and culturability

Although our initial screening yielded intriguing results, the critical downstream mechanisms underlying our observed phenomenon still remain elusive. Thus, we conducted a second screening using the *E. coli* BW25113 Keio knockout collection to assess the impact of UV on mutagenesis and culturability of the knockout strains. This screening focused on all potential repair mechanisms involved^16^, particularly (i) those that are not regulated by RecA (such as, MMR genes) and (ii) those that are regulated by RecA but whose expression was not significantly upregulated in our initial screening (**Fig. 2a**). For this second screening, the knockout strains in 96-well plates were treated with UV-B for 16 min during mid-exponential-phase growth, and RIF-resistant colonies were quantified after a 24-hour recovery period. We identified several genes whose deletion resulted in a significant reduction in RIF-resistant colony formation (**Fig. 3a**), including *recB*, a component of Exonuclease V or the RecBCD complex that promotes homologous recombination in the repair of double-strand DNA breaks^85,86^; *umuC* and *umuD*, encoding error-prone DNA polymerase V^87^; *ruvC*, encoding an endonuclease that binds to and cleaves Holliday junctions^88^; and *katE*, encoding *E. coli* catalase enzyme^89^. Additionally, we observed several genes that showed an increase in mutagenesis, such as *uvrD* and *uvrA*, which are involved in the nucleotide excision repair pathway^90,91^, and *mutY*, involved in the base excision repair pathway^92^ (**Fig. 3a**). Knockouts of MMR genes *(mutS, mutL, mutH,* and *exoI)*, which function in replication error correction, were included in our screen but did not show significant reductions in mutagenesis, indicating MMR may play a minimal role under our UV-B exposure conditions (**Fig. 3a**).

**Fig. 3:**
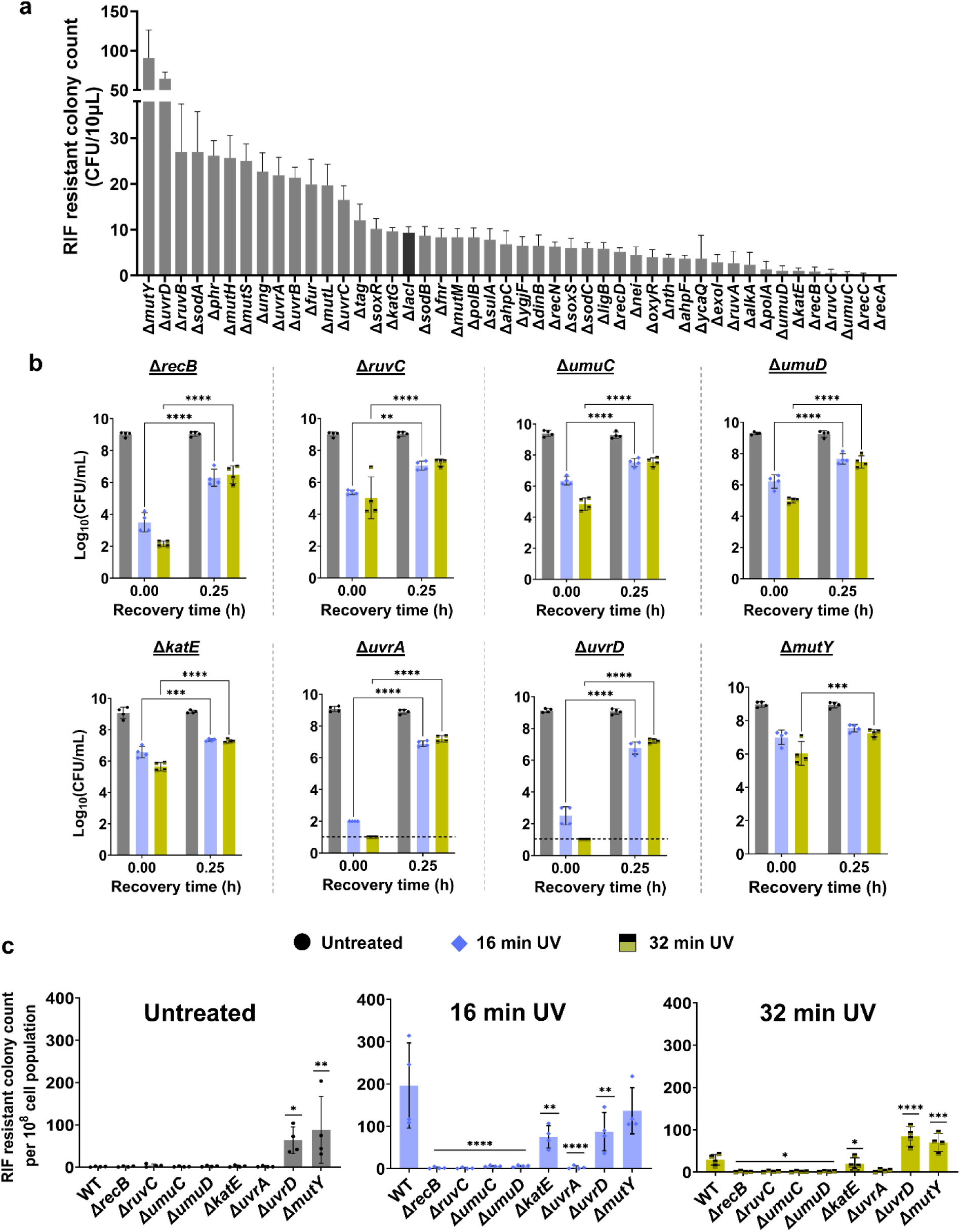
Knockout library screening to elucidate their effects on mutagenesis and cell culturability. (a) The *E. coli* BW25113 Keio knockout collection was screened following 16 min of UV treatment. RIF-resistant cells were quantified by spotting 10 μL samples from each well onto RIF-agar plates after a 24-hour recovery period. Δ*lacI* was used as the reference control (see Materials and Methods). (b) Selected genes were deleted in *E. coli* MG1655 and the mutagenesis assay was performed. Mid-exponential phase cells were exposed to UV for 0, 16 and 32 min and recovered for 24 h. At indicated time points (t = 0 h, 0.25 h, 0.5 h, 1 h, 2 h, 4 h, 6 h and 24 h) during recovery, cells were collected and plated to enumerate CFU. Note that only the 0 and 0.25 h time points are shown here; the full-time course is provided in **Supplementary Fig. S4.** (c) RIF-resistant cells were quantified by plating the samples on RIF-agar plates after 24 h of recovery. The values were reported as RIF-resistant colony count per 10^8^ cell population for different UV exposure durations. n=4. Statistical analysis was performed using one-way ANOVA with Dunnett’s post-test, where \**P* < 0.05, \*\**P* < 0.001, \*\*\**P* < 0.01. Data corresponding to each time point represents mean value ± standard deviation.

To further investigate genes that showed the most pronounced impact on UV-induced mutagenesis in our screening, and to see if similar trends occur in other *E. coli* strains, we deleted these genes in *E. coli* MG1655 and analyzed the knockout strains after 16 and 32 min of UV-B exposure. Deletion of *recB*, *ruvC*, *umuC*, and *umuD* significantly reduced mutagenesis (**Fig. 3c**), consistent with our screening data. The absence of UmuC and UmuD proteins, specialized for translesion synthesis, was shown to sensitize *E. coli* cells to UV^93,94^, aligning with our findings. While Δ*uvrA* in the *E. coli* BW25113 strain from screening showed higher mutant levels compared to wild-type cells, its deletion in the MG1655 background resulted in a significant reduction in mutant formation (**Fig. 3c**), suggesting that the role of certain genes in mutagenesis can vary depending on the genetic background of the *E. coli* strain. Also, our results indicate that KatE, UvrD, and MutY may not be directly involved in the mutagenic response to UV radiation under the conditions tested here, as their deletions did not clearly impact UV-B-induced mutagenesis in *E. coli* MG1655 (**Fig. 3c**). Interestingly, Δ*uvrD* and Δ*mutY* strains exhibited increased RIF-resistant colonies in cultures not treated with UV-B (**Fig. 3c**). UvrD is involved in various DNA repair pathways, including NER and MMR processes^41,95,96^. MutY is a glycosylase enzyme that corrects adenine mismatches resulting from DNA replication errors, primarily in the BER pathway^92^. While the specific roles of UvrD and MutY in UV-B-induced mutagenesis are not evident in this study, the increased RIF-resistant colonies in untreated cultures might highlight their potential role in maintaining genomic stability under normal growth conditions, possibly by mitigating spontaneous mutations.

The temporal CFU profiles revealed interesting trends as well. Δ*ruvC*, Δ*umuC*, and Δ*umuD* strains showed no significant change in transient non-culturability immediately after UV-B treatment compared to the wild type (**Fig. 3b, Supplementary Fig. S4**). However, Δ*recB* exhibited a notable decrease in CFU levels, while Δ*uvrA* and Δ*uvrD* mutants showed even more drastic reductions following UV-B treatment compared to the wild type (**Fig. 3c**). This reduction in CFU levels was transient, as their levels increased approximately 10^7^-fold within 15 min of recovery. UvrA and UvrD together form key components of the NER pathway^95^. UvrA forms a complex with UvrB to detect DNA lesions, while UvrD functions as a helicase to unwind and remove damaged DNA segments^96,97^. As both UvrA and UvrD are regulated by the RecA protein^32^, it is not surprising that the deletion of *recA* resulted in drastic reductions in CFU levels following UV treatment (**Fig. 1f**). Since RecA is essential for inducing the SOS response, its absence means that many DNA repair genes, including *uvrA* and *uvrD*, are not upregulated in response to UV-induced DNA damage.

### Excessive UV exposure impairs cellular translation processes

Although our results highlight specific SOS proteins involved in mutagenesis and cell culturability, it remains unclear why excessive UV exposure impairs *recA* expression and, consequently, the SOS response. This impairment could be due to the activation of cellular mechanisms, such as growth arrest proteins, which may help limit excessive DNA damage and mutagenesis. Additionally, the production of toxic molecules, such as reactive oxygen species, or the inhibition of certain cellular functions, like translation, may also contribute to this effect. These mechanisms—namely the involvement of growth arrest proteins, reactive oxygen species, or the inhibition of translation—will be thoroughly explored in the subsequent section of our study.

Within the SOS response, two genes, *sulA* and *tisB*, play key roles in cell growth inhibition. The *sulA* gene in *E. coli* encodes a cell-division inhibitor that temporarily halts the cell cycle during the SOS response, preventing damaged DNA from prematurely segregating into daughter cells^98^. On the other hand, *tisB* encodes a toxin protein that reduces cellular ATP levels^50,99^. This protein may also induce reversible dormancy by suppressing cell metabolism. To investigate the individual and combined effects of these genes, we constructed knockout strains of *E. coli* MG1655 Δ*sulA*, Δ*tisB*, and Δ*sulA*Δ*tisB*. Similarly, these strains, at the mid-exponential phase, were exposed to varying durations of UV-B radiation followed by a 24-hour recovery period. *E. coli* MG1655 WT cells were used as the control group for comparison. Deleting *sulA* and *tisB* individually did not alter the trend of transient non-culturability observed in our experiments. Specifically, the CFU profiles initially showed a significant decrease after prolonged UV-B exposure of 24 and 32 min, followed by a sharp increase within the first 15 min of recovery, similar to the wild-type strain (**Fig. 1b vs Fig. 4a-c**). However, CFU levels of Δ*sulA*Δ*tisB* immediately after 24 and 32 min of UV-B exposure (corresponding to the t~0 point on the CFU plots) were nearly 10-fold higher compared to those of the wild-type or single knockout strains (**Fig. 1b vs Fig. 4a-c**). We observed approximately a 10-fold increase in CFU levels within the first 15 min of recovery in the double knockout strain, although this increase was not as pronounced as in the control or single knockout strains (**Fig. 4a-c, Supplementary Fig. S5**). While deleting both genes enhanced the culturability of cells following UV-B treatment, the phenomenon observed in the wild-type strain was not completely eliminated in the double knockout strain, highlighting the complexity of the underlying mechanisms. No distinct profiles of RIF-resistant mutants were observed after 24 hours of recovery in either single or double-knockout strains compared to the wild-type strain (**Fig. 1c vs Fig. 4d-f**), indicating that these genes are not directly involved in mutagenesis, although they may affect cellular culturability to some extent.

**Fig. 4:**
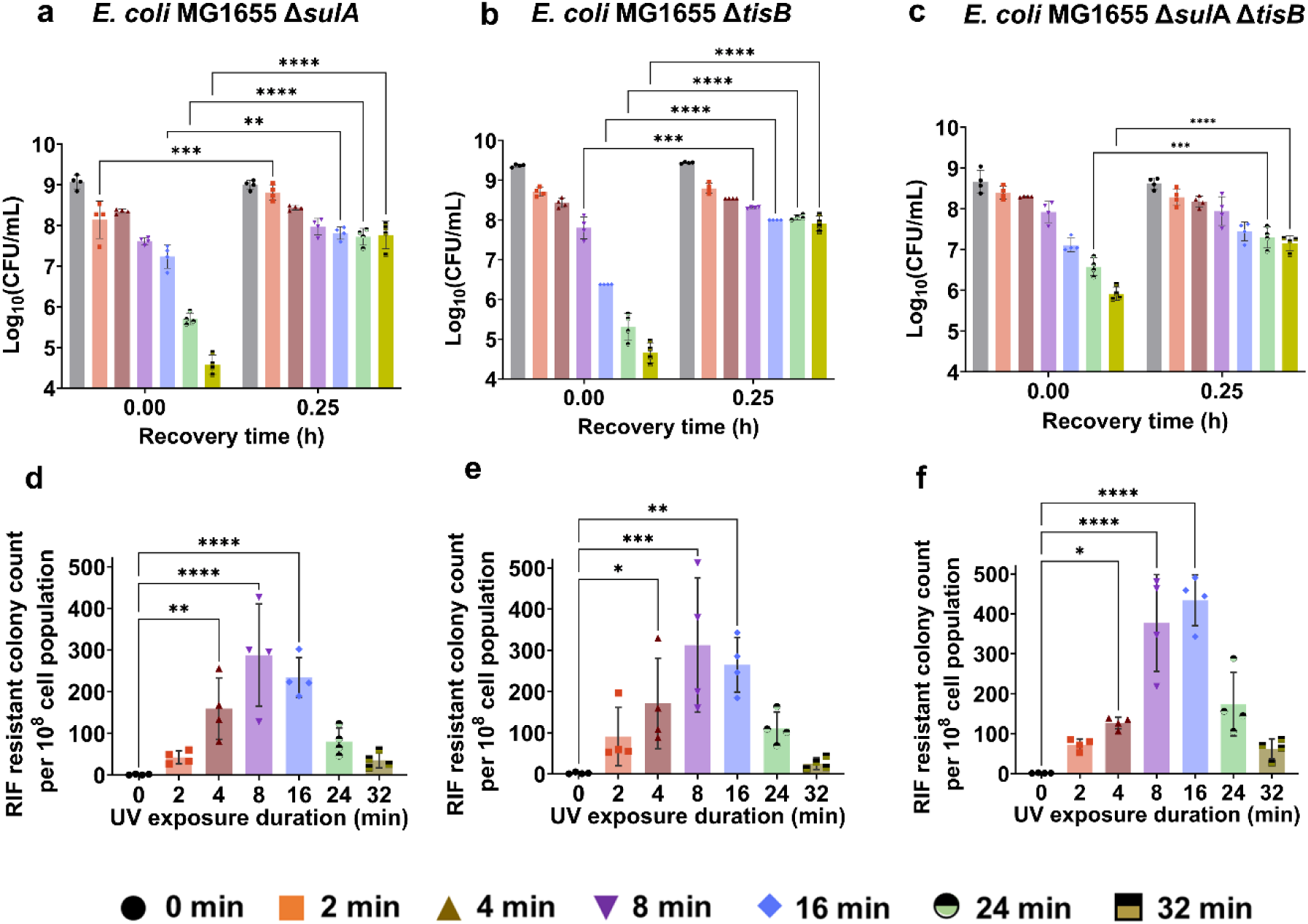
Perturbation of both SulA and TisB proteins moderately increased cell culturability during recovery. (a-c) Exponential-phase *E. coli* MG1655 Δ*sulA*, Δ*tisB*, and Δ*sulA*Δ*tisB* cells were exposed to UV-B light for 0, 2, 4, 8, 16, 24, and 32 minutes, followed by a 24-h recovery period. At specific time points during recovery, cells were collected and plated to determine their CFU levels. Note that only the 0 and 0.25 h time points are shown here; the full-time course is provided in **Supplementary Fig. S5.** (d-f) Levels of UV-induced RIF resistance mutations were measured by counting RIF-resistant colonies (per 10^8^ cells) in the cultures of three knockout strains after recovery for the indicated UV exposure times. n=4. Statistical analysis was performed using one-way ANOVA with Dunnett’s post-test, where \**P* < 0.05, \*\**P* < 0.001, \*\*\**P* < 0.01, \*\*\*\**P* < 0.0001. Data corresponding to each time point represents mean value ± standard deviation.

UV radiation has the potential to damage cellular membranes and compromise essential cellular functions^19^. This damage can lead to increased membrane permeability, leakage of cellular contents, and, ultimately, cell death^100,101^. Additionally, prolonged exposure to UV radiation can trigger excessive production of reactive oxygen species (ROS) within cells^102–104^. ROS are highly reactive molecules that can cause oxidative damage to cellular components such as DNA, proteins, and lipids, impairing cellular functions. It is possible that the 24-minute and 32-minute UV-B treatments studied here may not be severe enough to kill the bacterial cells completely, but they are likely sufficient to hinder cell growth and mutagenic processes by disrupting vital metabolic processes. To characterize cellular membrane permeability, cells treated with UV-B were immediately collected and incubated with propidium iodide (PI), a fluorescent dye that enters cells with compromised membranes. Flow cytometry data indicated no significant increase in permeabilization following prolonged UV-B treatment (32 min) up to 1 hour of recovery (**Fig. 5a**), compared to both untreated cells and moderately UV-treated cells (16 min), which did not markedly reduce CFU levels. Furthermore, when UV-treated samples were subjected to Amplex Red reagent (see Materials and Methods for details) to detect hydrogen peroxide (H₂O₂), a primary reactive oxygen species, no significant changes were observed in the 32-minute UV-treated cells compared to untreated or 16-minute UV-treated cells (**Fig. 5b**), indicating H₂O₂ may not be a key player. In our previous study, we found that ROS significantly affects cell culturability only when their concentrations are much higher than physiological levels, and their impact on cell culturability was observed to be non-transient ^54^.

**Fig. 5:**
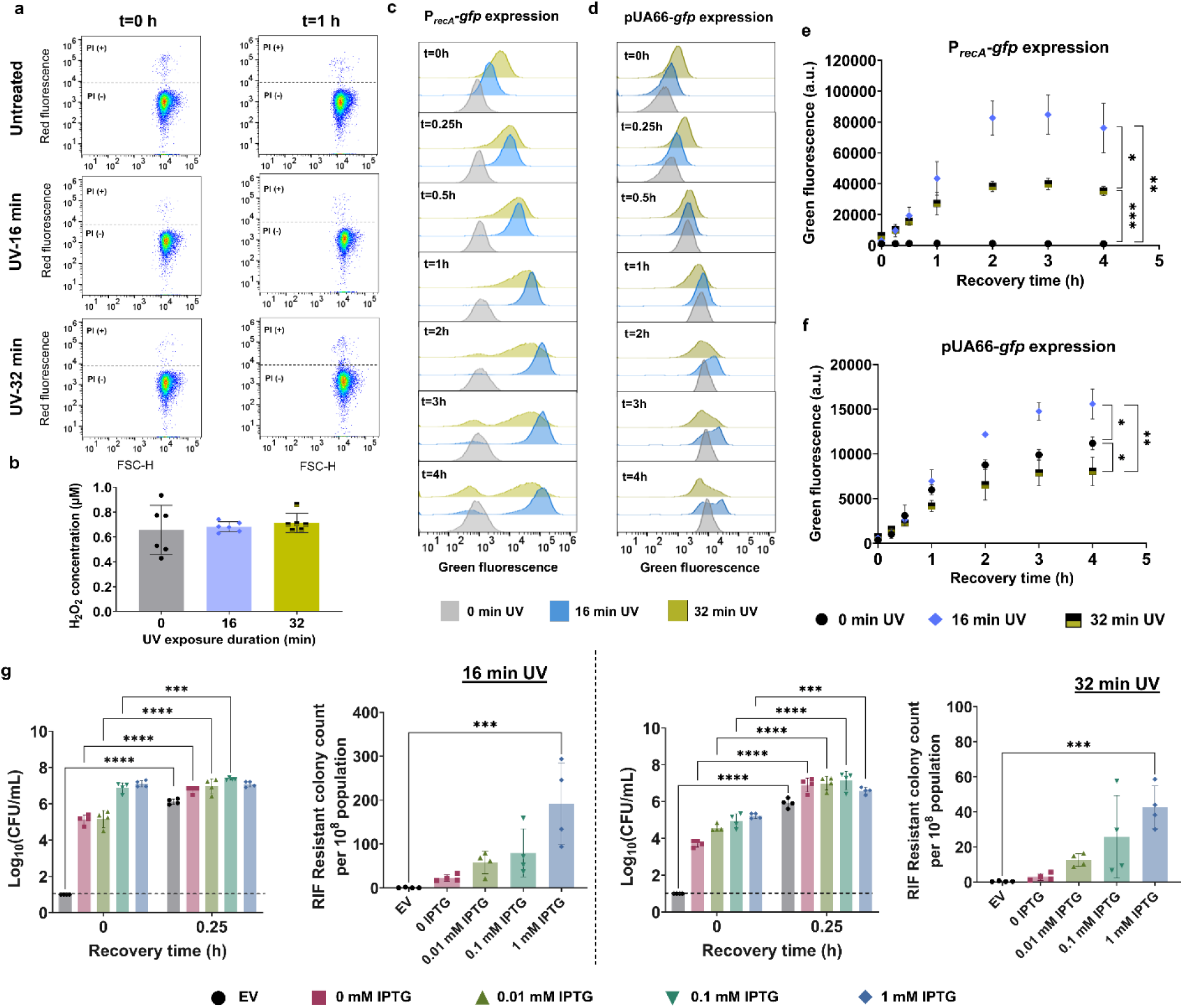
Excessive UV exposure did not compromise cell membrane integrity or increase hydrogen peroxide levels, but it did impair cellular translation. (a) Membrane integrity of UV-treated cells (0, 16, and 32 min of UV exposure) was assessed via flow cytometry with PI staining (see **Supplementary Fig. S8** for controls). FSC-H: Forward light scatter. (b) H_2_O_2_ concentrations were measured using the Amplex Red Hydrogen Peroxide/Peroxidase Assay Kit. Reaction mixtures containing Amplex Red reagent, horseradish peroxidase, and UV-treated cells in sodium phosphate buffer (pH 7.4) were incubated for 30 minutes at room temperature, followed by fluorescence measurement with a microplate reader. H_2_O_2_ levels were calculated using a standard curve (**Supplementary Fig. S9**). (c) Expression of P*_recA_*-*gfp* in UV-treated *E. coli* MG1655 cells was analyzed via flow cytometry for the indicated time points during recovery (P*_recA_* : the *recA* promoter). (d) IPTG-inducible GFP expression in UV-treated *E. coli* MG1655 pUA66-*gfp* cells was also assessed by flow cytometry for the indicated time points during recovery. (e) GFP expression profiles of UV-treated cells harboring the *recA* promoter and (f) IPTG-inducible GFP expression systems were evaluated using mean GFP values from flow cytometry data. (g) Recovery (CFU counts) and mutation frequency of *E. coli* MG1655 Δ*recA* carrying an IPTG-inducible *recA* overexpression plasmid were assessed following 16- and 32-minute UV-B exposures, shown for the indicated IPTG concentrations. n ≥ 4. Statistical analysis was performed using one-way ANOVA with Dunnett’s post-test, where \**P* < 0.05, \*\**P* < 0.001, \*\*\**P* < 0.01, \*\*\*\**P* < 0.0001. Data corresponding to each time point represent mean value ± standard deviation.

UV exposure is known to interfere with ribosomal function by inducing crosslinks in ribosomal RNAs^105^, which can inhibit overall translation processes and impair the expression of SOS response genes. To assess the effects of UV treatments on cellular translation, we analyzed GFP expression under the control of the *recA* promoter (P*_recA_*) at the single-cell level using *E. coli* MG1655 cells carrying the pUA66 P*_recA_*-*gfp* plasmids. The cells were subjected to UV-B treatments for 16 min and 32 min, and *recA* expression was measured using flow cytometry during the early recovery phase (**Fig. 5c**). UV-B treatment should activate promoters of SOS response genes and induce recA expression. As expected, the 16-min UV-B treatment resulted in significantly higher *recA* expression compared to untreated cells (**Fig. 5c, e**). However, the 32-min UV-B exposure resulted in more heterogeneous and lower *recA* expression compared to the 16-min UV-B exposure (**Fig. 5c, e**), potentially implying an impairment in the translational process. This difference between the 16- and 32-min treatments cannot be solely attributed to increased cell death following 32 min of UV-B exposure. Despite the observed transient culturability in the prolonged UV-exposure culture, CFU levels in both conditions are nearly similar after the brief transient period, around 15 minutes of recovery culture (see the 0.25-hour time point in **Fig. 1b**).

To substantiate the hypothesis regarding the impairment of the translational process, we employed a GFP-expressing plasmid where the GFP gene is tightly regulated by an isopropyl β-D-1-thiogalactopyranoside (IPTG)-inducible T5 promoter and a strong LacI^q^ repressor^106^. Our rationale was that if excessive UV-B treatment indeed impairs translational processes, the levels of GFP expression in cells treated with 32 min of UV-B should be lower compared to those in the no-UV control or 16-minute UV-treated cells, despite the presence of the inducer. We supplemented cell cultures with 0.1 mM IPTG immediately before UV-B treatments. As anticipated, the 32-min UV-B treatment resulted in a lower level of GFP expression compared to the no-UV control or the 16-minute UV-B treatment (**Fig. 5d, f**). However, the 16-min UV-B treatment exhibited the highest GFP expression levels (**Fig. 5d, f**), suggesting that moderate UV treatment (16 min) enhances translational activity, whereas excessive UV-B treatment (32 min) slightly inhibits it. This moderate inhibitory effect of excessive UV-B treatment on cellular translation, including RecA expression, likely impacts overall cellular processes and compromises the SOS response. This was further validated when we measured the expression levels of *uvrD* and *uvrA*—two critical genes involved in mutagenesis and culturability. Their expression levels were slightly reduced under 32 minutes of UV treatment compared to 16 minutes (**Supplementary Fig. S6a-d**).

To further validate the role of RecA-mediated SOS response in recovery and mutagenesis following UV-B exposure, particularly under conditions where transcription and translation are impaired, we constructed an IPTG-inducible *recA* expression system. This was achieved by cloning *recA* under the control of a T5 promoter (**Supplementary Fig. S7**) and introducing the plasmid into the *E. coli* MG1655 Δ*recA* background. In the absence of IPTG, the strain behaves as a *recA* knockout, showing impaired recovery and loss of UV-induced mutagenesis (**Fig. 5g**). To ensure that RecA protein was present at the time of UV treatment, IPTG was added approximately 30 minutes prior to exposure. This timing was chosen to avoid possible interference from UV-induced translation defects that could impair *recA* expression if induction occurred during or after treatment. Furthermore, we tested a range of IPTG concentrations to evaluate whether RecA-dependent outcomes were dose-responsive, as suggested by our data here and in our previous study ^54^. As shown in **Fig. 5g**, increasing IPTG levels led to a corresponding restoration of both CFU levels and mutation frequency following 16- and 32-minute UV-B exposures, indicating a clear correlation between RecA expression and the recovery of culturability and mutagenesis. These findings suggest that translational capacity directly impacts SOS activation by limiting RecA availability.

Altogether, our findings suggest that while prolonged UV-B treatments may not directly impact membrane integrity or ROS levels, the observed unculturability of the cells likely results from a combined effect of multiple factors, including cell growth inhibitors, impaired translation processes, and downstream repair pathways. The impairment of cellular processes such as translation may hinder the SOS response, as evidenced by the decreased expression of RecA in cells treated with 32 min of UV-B (**Fig. 5e**). This interpretation is further supported by our findings with the Δ*recA* strain, which exhibited near-complete unculturability immediately after UV treatments. The immediate reduction in CFU levels in the knockout strains (Δ*recA*, Δ*uvrA* and Δ*uvrD*), as well as in the wild-type strain exposed to prolonged UV, highlights the importance of the SOS response in the initial phases of DNA damage repair. The transient nature of CFU reduction in all these mutants also suggests that *E. coli* possesses redundant or numerous repair mechanisms^16^ that can partially restore culturability even when certain DNA repair pathways are disrupted.

## Conclusions

In this study, we investigated the effects of UV-B treatments on *E. coli* mutagenesis and culturability and provided important insights into bacterial survival mechanisms under stress. Mutagenesis, measured by RIF-resistant colony formation, peaked at 16 min of UV-B exposure but decreased with longer exposures (24 and 32 min). The longer UV-B exposures also led to a significant decrease in CFU levels, but this phenomenon was observed to be a transient state, as we saw a drastic jump in CFU levels after a short recovery period. We also explored the role of key SOS response genes and demonstrated that knockout strains lacking *sulA* and *tisB* genes exhibited similar patterns of transient non-culturability as wild-type cells, although double knockouts exhibited higher initial CFU levels post-exposure, indicating a partial role in culturability. Further investigation revealed that excessive UV exposure did not significantly affect membrane permeability or reactive oxygen species (ROS) levels but moderately impaired translation processes, as evidenced by reduced expression of SOS response genes like *recA* or IPTG-inducible fluorescent proteins. This impaired translation likely contributed to the observed reduction in mutagenesis and transient non-culturability following prolonged UV exposure, given the significant impact of *recA* deletion on cell culturability and mutagenesis. Importantly, reintroducing *recA* expression in the knockout strain restored culturability and increased mutagenesis in proportion to *recA* expression levels, further supporting the link between translational capacity, *recA*-mediated DNA repair, and the observed phenotypes.

Although our screening of promoter reporters highlighted several genes significantly upregulated following UV treatment, the knockout strains of most of these genes generally had minimal impact on culturability or mutagenesis, except for Δ*recN* and Δ*rmuC*, which moderately reduced mutagenesis. In our second screening with the Keio knockout collection, we identified several critical genes, such as *recB, umuC, umuD*, and *ruvC*, whose deletions led to decreased mutagenesis, highlighting their crucial roles in facilitating error-prone repair mechanisms during UV-B stress. Furthermore, we observed differential impacts of gene deletions on mutagenesis versus culturability, providing insights into additional functions of these genes. Particularly in strains lacking *uvrA*, or *uvrD* showed a drastic but transient reduction in their culturability following UV treatment, highlighting their role in maintaining genomic stability. Our findings also underline the abundance of repair mechanisms in *E. coli*, demonstrating that multiple independent pathways can significantly impact cell mutagenesis and culturability.

Overall, this study enhances our understanding of bacterial responses to environmental stressors, particularly UV-B radiation, by uncovering the complex interplay between DNA repair pathways, mutagenesis, and transient non-culturability. Our findings reveal a dose-dependent balance between survival and genetic variability, suggesting that bacteria modulate their repair and stress responses to optimize adaptability under UV-B exposure. This regulatory balance has important implications for understanding how UV-B, especially during periods of ozone depletion, may act as a powerful evolutionary force.

## Materials and Methods

### Bacterial strains and plasmids

*Escherichia coli* K-12 MG1655 wild type (WT), the pUA66 plasmid containing an IPTG-inducible *gfp* (green fluorescent protein gene) expression system, and the pUA66 plasmid with *gfp* under the control of the *recA* promoter, P*_recA_*, were obtained from Dr. Mark P. Brynildsen at Princeton University. *E. coli* K-12 MG1655 Δ*sulA,* Δ*tisB,* Δ*recA* was constructed in our previous studies ^54,107,108^. The promoter strain collection of *E. coli* MG1655 in a 96-well plate format used for the screening assay was obtained from Horizon Discovery, Lafayette, CO, USA. The Keio knockout strain collection (a single-gene deletion library of *E. coli* K-12 BW25113) was obtained from Dharmacon Keio Collection (Dharmacon, Cat# OEC4988). Since the knockout strains carry a kanamycin resistance gene, high-throughput screening was performed in the presence of kanamycin to prevent contamination; thus, Δ*lacI* was used as the reference control, given that the parental BW25113 strain lacks kanamycin resistance. We selected the Δ*lacI* mutant as it is not involved in DNA repair pathways and serves as a neutral background. Furthermore, its response to UV-B exposure closely mirrors that of the MG1655 wild-type strain (**Supplementary Fig. S2a**), supporting its suitability as a functional reference for our experimental comparisons. The strains from both the promoter and knockout collections used in this study are listed in **Supplementary Table 1**. The knockout *E. coli* MG1655 strains generated for this study are detailed in **Supplementary Table 2**. The method of Datsenko and Wanner^69^ was used to generate these strains and the oligonucleotides used to delete the genes are provided in **Supplementary Table 3.** We also attempted to generate a Δ*phr* mutant in *E. coli* MG1655 using the same approach; however, no viable recombinants were obtained despite testing multiple primer sets with varying homology arm lengths (40–60 bp) (**Supplementary Table 3**). An IPTG-inducible *recA* overexpression plasmid was generated using a commercial cloning service (Synbio Technologies, USA). The *recA* coding sequence from *E. coli* MG1655 was cloned downstream of a T5 promoter into a low-copy plasmid backbone (pUA66) containing a strong mutated *lacI* repressor for tight regulation and a kanamycin resistance marker^108^. The resulting plasmid was verified by sequencing and subsequently transformed into MG1655 Δ*recA* cells for inducible expression studies. The plasmid map is shown in **Supplementary Fig. S7.**

### Chemicals, media, and culture conditions

Unless otherwise specified, all chemicals were procured from Fisher Scientific (Atlanta, GA), VWR International (Pittsburgh, PA), or Sigma Aldrich (St. Louis, MO). PI staining kit was purchased from Promega Corporation (Madison, WI). *E. coli* cells were cultured in liquid Lysogeny-Broth (LB) medium. LB agar medium was utilized for enumerating colony-forming units (CFU) of *E. coli*. The liquid LB medium was prepared by dissolving 5 g yeast extract, 10 g tryptone, and 10 g sodium chloride in 1 L of deionized (DI) water. LB Agar media were prepared by dissolving pre-mixed 40 g LB agar in 1 L of DI water. Both solid and liquid media were subjected to autoclaving for sterilization.

Kanamycin (50 µg/mL) was included in the liquid LB media for plasmid selection and retention. IPTG at 0.1 mM was used to induce *gfp* expression. For PI staining, sterile 0.85% sodium chloride solution was used. When necessary, cells were washed with phosphate-buffered saline (PBS, 1X). Stock solutions for rifampicin (RIF; 50 mg/mL) was prepared by dissolving in DI water using 0.01 N sodium hydroxide. The IPTG stock solution (1 mM) was dissolved in DI water. All chemical solutions were sterilized using 0.2 μm VWR syringe filters.

To prepare RIF-agar plates, the stock RIF solution was added to autoclaved LB agar, resulting in a final plate concentration of 500 μg/mL RIF. Unless specified otherwise, overnight pre-cultures were generated in 14-mL Falcon test tubes containing 2 mL of liquid media. These pre-cultures were inoculated from a 25% glycerol cell stocks stored at −80 °C and cultivated for 24 h at 37 °C with shaking at 250 revolutions per min (rpm). Experimental cell cultures were prepared by diluting the overnight pre-cultures (1:100) into 2 mL of fresh LB medium in 14-mL Falcon test tubes. Bacterial cells in this study reached the mid-exponential phase (OD_600_ ~ 0.5) after around 3 h, attaining an average cell density of 7×10^8^ CFUs/ml. All treatments involving UV were administered at this stage.

### UV treatment and cell recovery

Overnight pre-cultures of *E. coli* MG1655 cells were diluted 100-fold in 2 mL fresh LB media in test tubes and grown at 37°C with shaking (250 rpm). Cell growth was monitored by measuring optical density at 600 nm wavelength (OD_600_) with a plate reader (Varioskan LUX Multimode Microplate Reader, Thermo Fisher, Waltham, MA, United States). When the cell density reached OD_600_ ~ 0.5, cultures from the test tubes were transferred to petri dishes (the diameter of petri dishes= 100 mm, catalog no. FB0875713, Fisher Scientific). This procedure created a thin film of culture in the dish with a height of about 0.25 mm. This configuration resulted in a uniformly distributed liquid film without observable surface tension-driven droplet formation (i.e., beading). The cultures were exposed to a light source emitting UV-B at 302 nm (UVP ChemStudio, catalog no. 849-97-0928-02; Analytik Jena, Jena, Germany) for varying exposure times (0, 2, 4, 8, 16, 24 and 32 min). In this setup, UV-B light was emitted from below using a transilluminator and passed through a UV-permeable plastic Petri dish placed directly on the transilluminator surface. This configuration ensured uniform and efficient bottom-up exposure of bacterial cells within the thin culture film. To verify the consistency and intensity of UV-B irradiation, direct irradiance measurements were performed using a Vernier™ UV-B Sensor (Fisher Scientific, Cat. No. S16273ND), calibrated for peak sensitivity at 315 nm (detection range: 290–320 nm), in combination with a LabQuest™ 3 data acquisition platform. Measurements were taken under four conditions to assess the impact of the experimental setup on UV transmission:(a) sensor placed directly on the transilluminator surface without a Petri dish, (b) sensor on an empty Petri dish, (c) sensor inside a Petri dish containing LB medium (not in contact with the liquid), and (d) sensor in direct contact with the LB medium inside the dish. In all cases, UV-B irradiance was consistently measured at approximately 1000 mW/m^2^ (equivalent to 1 W/m^2^), indicating that neither the Petri dish nor the medium significantly interfered with UV transmission (see **Supplementary Fig. S1**). The energy dose (in J/m^2^) for each exposure condition was calculated using the formula:

### Energy dose (J/m^2^) = Irradiance (W/m^2^) × Exposure time (s)

After UV exposure for the indicated time points, cells with the LB media were transferred back to test tubes and recovered for 24 h. During the recovery period, 10 μl samples were collected at specified time points from each test tube, serially diluted in PBS in round-bottom 96-well plates, and then, plated on LB agar media. The plates were incubated at 37 °C for 16 h to enumerate CFU. We note that new colonies were not formed when incubated beyond 16 h. A similar procedure was followed for the other strains. For dark recovery experiments, all procedures prior to UV treatment were identical to those described above. Following UV exposure, samples were incubated in a large, dark incubator-shaker, with the front window fully covered in aluminum foil to prevent light exposure and photoreactivation. Additionally, the test tubes containing the cultured cells were completely wrapped in aluminum foil to ensure total darkness during the recovery period.

### Assessing mutagenesis

To assess the extent of mutant cell formation induced by UV exposure, cells were exposed to UV radiation for the specified durations, followed by a 24-h recovery period. After this recovery period, 500 μL of cells were collected and spread onto agar plates containing 500 μg/mL of RIF. These plates were then incubated at 37 °C for 16 h to enumerate RIF-resistant colonies. To determine clonogenic survival, CFU levels were determined, and mutant levels were normalized by dividing the number of RIF-resistant colonies by the total number of colonies in a 1 mL culture volume. Unless specified otherwise, mutant formation was reported as the count of RIF-resistant colonies per 10^8^ cell population.

### PI Staining

Mid-exponential *E. coli* cells were treated with UV for 16 min and 32 min duration following the protocol mentioned above. At the beginning of the recovery and 1h after the recovery (t=0 h and 1 h), UV-treated cells in LB media were diluted 20-fold in 1.0 ml 0.85% NaCl solution in flow cytometry tubes (5 ml round bottom Falcon tubes, size: 12 × 75 mm) to achieve the desired cell density (~10^6^-10^7^ cells/ml) for flow cytometry analysis. The resulting cell suspensions were treated with 20 µM PI dye. PI produces red fluorescence upon binding DNA; however, it can only penetrate cells with damaged membranes. The samples were incubated in the dark at 37 °C for 15 min before analyzing them with a conventional bench-top flow cytometer (NovoCyte 3000RYB, ACEA Biosciences Inc., San Diego, CA, United States). For flow cytometry analysis, we chose a slow sample flow rate (14 µl/ min) to have a sample stream diameter (i.e., core diameter) of 7.7 µm. The instrument has a constant sheath flow rate of 6.5 ml/min. The flow cytometer utilizes low-power solid-state lasers. Cells were excited at a 561 nm wavelength and red fluorescence was detected with a 615/20-nm bandpass filter. At least 30,000 events were recorded for each sample. NovoExpress software was used to collect the data. PI-stained dead cells, obtained after ethanol (70% v/v) treatment, were used as a positive control. PI-stained live cells were used as a negative control. Forward and side scatter signals of untreated live cells were used to determine the cells on flow cytometry diagrams; the positive and negative controls were used to gate PI positive (+) and PI negative (-) cell populations **(Supplementary Fig. S8).**

### H_2_O_2_ measurement

The Amplex™ Red Hydrogen Peroxide/Peroxidase Assay Kit (Invitrogen, Thermo fischer scientific, Catalog number: A22188) was used to assess the amount of H_2_O_2_ formed in the UV-treated cells. The Amplex™ Red Hydrogen Peroxide/Peroxidase Assay Kit contains a sensitive, one-step assay that uses the Amplex™ Red reagent (10-acetyl-3,7-dihydroxyphenoxazine) in combination with horseradish peroxidase (HRP) to detect hydrogen peroxide (H_2_O_2_). In the presence of peroxidase, the Amplex™ Red reagent reacts with H_2_O_2_ in a 1:1 stoichiometry to produce the red-fluorescent oxidation product, resorufin. Resorufin has excitation and emission maxima of approximately 571 nm and 585 nm. For preparing the stock solutions, 10 mM Amplex® Red reagent was prepared by dissolving the contents of the vial of Amplex® Red reagent in 60 μL of DMSO. Reaction buffer was diluted 5-fold to prepare 1X reaction buffer and 10 U/mL Horseradish Peroxidase (HRP) was prepared by dissolving one vial of HRP in 1.0 mL of 1X Reaction Buffer. The working solution of 100 μM Amplex® Red reagent and 0.2 U/mL HRP was prepared by adding 50 μL of 10 mM Amplex® Red reagent stock solution and 100 μL of 10 U/mL HRP stock solution in 4.85 mL of 1X Reaction Buffer. The assay volume for this experiment was 100 μL. Mid-exponential *E. coli* cells were treated with UV for 16 min and 32 min duration and right after the treatment, 50 μL of sample for each condition was serially diluted in 1X Reaction Buffer to determine the optimal amount of sample for the assay. 50 μL of the Amplex® Red reagent/HRP working solution was added to each microplate well containing the samples. The samples were incubated at room temperature for 30 min, and protected from light. At the end, the fluorescence was measured using a microplate reader equipped for excitation in the range of 530– 560 nm and fluorescence emission detection at ~590 nm. For the standard curve, 20 mM Hydrogen Peroxide (H_2_O_2_) working solution was prepared by dissolving 3.0% (0.88 M) H_2_O_2_ in 1X Reaction Buffer and then further diluting 20 mM H_2_O_2_ working solution into 1X Reaction Buffer to produce H_2_O_2_ concentrations of 0 to 10 μM, each in a volume of 50 μL. 50 μL of the Amplex® Red reagent/HRP working solution was added to each microplate well containing the standards followed by 30 min of incubation and then measuring the fluorescence following the same way **(Supplementary Fig. S9).**

### Fluorescent protein expression assay for reporter genes

Overnight pre-cultures of *E. coli* MG1655 cells with reporter genes fused to the SOS gene promoters (P*_recA_*, P*_uvrA_*, and P*_uvrD_*) were diluted 1:100 in 2 mL of LB media within test tubes and incubated at 37°C with shaking (250 rpm). Upon reaching the mid-exponential phase, the cells were subjected to UV radiation for specified durations (16 min and 32 min). An untreated culture was used as a control. At the beginning of the recovery, 20 µM PI dye was added to the cultures for continuous monitoring of the membrane permeability. At specified time points during the recovery (t=0, 0.25, 0.5, 1, 2, 3 and 4 h), UV-treated cells were diluted 20-fold in 1.0 ml 0.85% NaCl solution in flow cytometry tubes (5 ml round bottom Falcon tubes, size: 12 × 75 mm) to achieve the desired cell density (~10^6^-10^7^ cells/ml). The samples were analyzed using the same flow cytometry method described above (see the “PI staining” section); however, cells were analyzed using two lasers. For measuring the red fluorescence from PI dye, cells were excited at a 561 nm wavelength, and red fluorescence was detected with a 615/20-nm bandpass filter. For measuring the green fluorescence, cells were excited at a 488 nm wavelength, and the green fluorescence was detected with a 530/30 bandpass filter.

### Transcription/translation activities of pUA66-*gfp* plasmid

To assess the effect of the UV treatment on transcription/translation, the amount of GFP produced by *E. coli* strains from the low-copy plasmid was measured. The plasmid, pUA66-*gfp*, has a *gfp* gene under the control of a strong, IPTG-inducible T5 promoter and a strong LacI^q^ repressor. Overnight pre-cultures of *E. coli* MG1655 cells carrying pUA66-*gfp* were diluted 100-fold in 2 mL fresh LB media in test tubes and grown at 37°C with shaking (250 rpm). At the mid-exponential phase (OD_600_~0.5), 0.1mM IPTG was added to the test tubes and then immediately treated with UV for 16 min and 32 min duration. Untreated cultures having IPTG only (without UV exposure) served as controls. At specified time points during the recovery (t=0, 0.25, 0.5, 1, 2, 3, and 4 h), UV-treated cells were diluted 20-fold in 1.0 ml PBS in flow cytometry tubes (5 ml round bottom Falcon tubes, size: 12 × 75 mm) to achieve the desired cell density (~10^6^-10^7^ cells/ml). The samples were analyzed using the same flow cytometry method described above (see the “PI staining” section); however, cells were analyzed with a laser emitting light at 488 nm and the green fluorescence was detected with a 530/30 bandpass filter.

### Promoter library screening

Overnight precultures were prepared by inoculating the strains from the promoter library into the wells of 96-well plates containing 200 μl LB medium and 50 µg/ml kanamycin (for plasmid retention). The plates were sealed with a sterile, oxygen-permeable membrane (Breathe-Easier, Cat# BERM-2000, VWR International) and cultured for 24 h at 37°C with shaking at 250 rpm. Overnight precultures were diluted 1:40 in LB medium with kanamycin in a new 96-well plate, sealed, and incubated at 37°C with shaking at 250 rpm. At the mid-exponential phase (OD_600_ = 0.5), cultures in 96 well plates were exposed to UV light (UVP ChemStudio, catalog no. 849-97-0928-02; Analytik Jena, Jena, Germany) for 16 min, and then, recovered for 24 h at 37°C with shaking at 250 rpm. GFP was measured with a Varioskan LUX Multimode Microplate Reader (Thermo Fisher, Waltham, MA, USA) at the indicated times with untreated cultures as a control. The excitation and emission wavelengths for GFP measurement were 485 nm and 511 nm, respectively. Fold changes for GFP (treated/untreated cultures after 24 h treatment) were used to report the promoter activity. The top 10 promoters showing the highest expression were selected for subsequent experiments.

### Screening *E. coli* BW25113 Keio knockout collection

Overnight cultures of single mutants **(Supplementary Table 1)** carrying kanamycin resistance genes were diluted 40-fold into flat-bottom 96-well plates, with each well containing 200 μL of cell culture. Kanamycin (50 µg/ml) was added to overnight and treatment cultures to prevent contamination. The plates were securely sealed using a sterile, oxygen-permeable membrane (Breathe-Easier, Cat# BERM-2000, VWR International) and incubated at 37°C with agitation at 250 rpm. When the cultures reached the mid-exponential phase (OD_600_ ~ 0.5), the cells in the 96-well plates were exposed to UV light (UVP ChemStudio, catalog no. 849-97-0928-02; Analytik Jena, Jena, Germany) for 16 min. Subsequently, they were allowed to recover for 24 h at 37°C with shaking at 250 rpm. To quantify UV-induced mutant cells, 10 μL samples were collected from each well and spotted onto LB agar plates, containing 500 μg/mL RIF, at the end of the 24-h recovery period. The plates were then incubated at 37°C for 16 h to enumerate RIF-resistant colonies. The mutant formation was reported as the count of RIF-resistant colonies per 10 μL of cell culture.

### Overexpression of *recA* Using an IPTG-Inducible Plasmid

Overnight pre-cultures of *E. coli* MG1655 Δ*recA* cells harboring the pUA66-*recA* plasmid were diluted 1:100 into 2 mL of fresh LB medium in test tubes and incubated at 37 °C with shaking at 250 rpm. After 2.5 hours of growth, IPTG was added to final concentrations of 0.01, 0.1, and 1 mM. One tube was maintained without IPTG as a non-induced control. Cultures were then incubated for an additional 30 minutes to allow *recA* expression prior to UV exposure. Subsequently, the cultures were exposed to UV for either 16 or 32 minutes. As a control, *E. coli* MG1655 Δ*recA* cells carrying the pUA66 empty vector were included for both UV exposure durations. During the recovery period, 10 μL samples were collected at defined time points from each culture, serially diluted in PBS using round-bottom 96-well plates, and plated on LB agar to determine CFU. Plates were incubated at 37 °C for 16 hours. RIF-resistant colonies were enumerated following the same procedure.

### Statistical analysis and reproducibility

For all pairwise comparisons, one-way ANOVA with Dunnett’s post-test was utilized. A minimum of four independent biological replicates (unless otherwise specified) were conducted for experiments involving UV exposure. In all figures, data corresponding to each time point represents the mean value ± standard deviation. Regarding statistical significance analysis, the threshold values were set as follows: *P < 0.05, **P < 0.01, ***P < 0.001, and ****P < 0.0001. All figures were created using GraphPad Prism 10.0.2, and the statistical analyses were carried out using GraphPad Prism 10.0.2 statistical functions. FlowJo V 10.7.1 was used to analyze the data obtained from flow cytometry.

## Acknowledgment

The authors would like to thank the members of Orman Lab for their help. This study was supported by NIH/NIAID R01-AI143643.

## Contributions

S.G. and M.A.O. conceived and designed the study. S.G., and J.N. performed the experiments.

S.G. and M.A.O. analyzed the data and wrote the paper. All authors read and approved the paper.

## Corresponding author

Correspondence to Mehmet A. Orman

## Declaration of interests

The authors declare no competing interests.

## Data availability

All data presented in this manuscript are available in the Main Text or the Supplementary File. All raw data have been published on Figshare: https://doi.org/10.6084/m9.figshare.27701493.v2

## Notes

### Competing Interest Statement

The authors have declared no competing interest.

### Summary of Updates

This revised version of the manuscript includes additional control experiments, such as dark controls and tests evaluating the effects of Phr on UV-B-induced mutagenesis, as well as the impact of an inducible recA overexpression system on UV-B-mediated transient culturability. Due to Jenet Narzary's significant contributions to these additional experiments, her name has been included as a co-author.

