## Supplemental Figures and Tables for "UV-B-Induced DNA Repair Mechanisms and Their Effects on Mutagenesis and Culturability in *Escherichia coli*"

**Sreyashi Ghosh[1], Jenet Narzary, and Mehmet A. Orman[1]***
[1]William A. Brookshire Department of Chemical and Biomolecular Engineering, University of Houston, Houston, TX, USA

### Supplementary figures

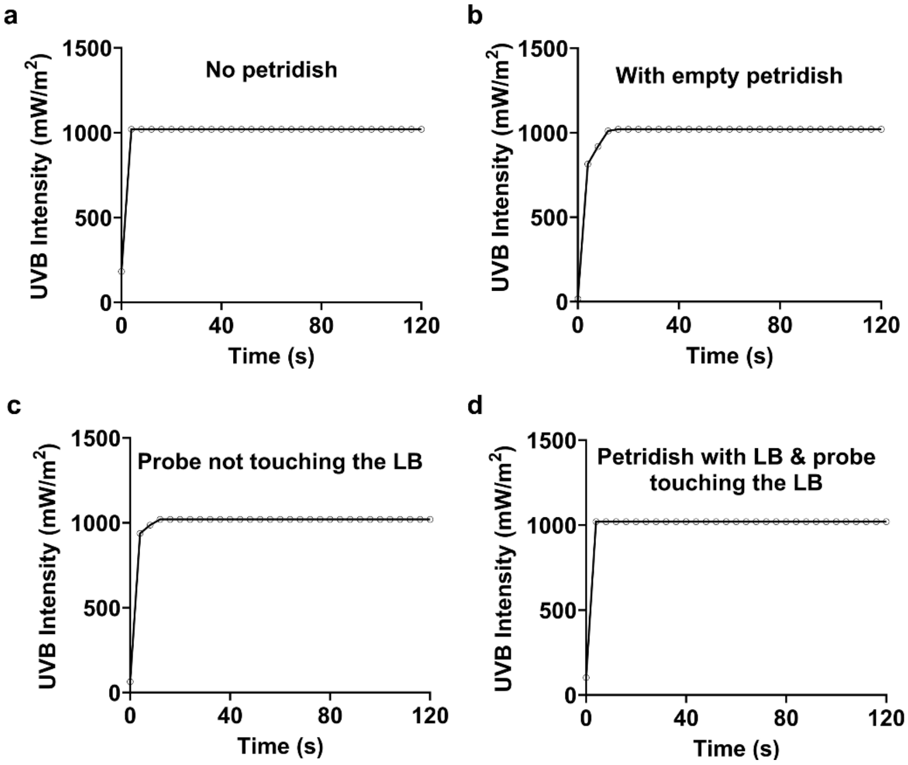

**Fig. S1. UV lamp intensity measurement at 302 nm using a UV-B dosimeter.** UV-B intensity was measured using a dosimeter (see Materials and Methods for specifications) under four different conditions to ensure accurate assessment of the irradiance reaching bacterial cells. (a) Probe placed directly on the surface of the UV transilluminator block with no Petri dish. (b) Probe placed on an empty Petri dish positioned on the UV transilluminator. (c) Probe positioned inside a Petri dish containing LB medium, without contacting the liquid surface. (d) Probe in direct contact with the LB medium inside the Petri dish on the UV transilluminator block.

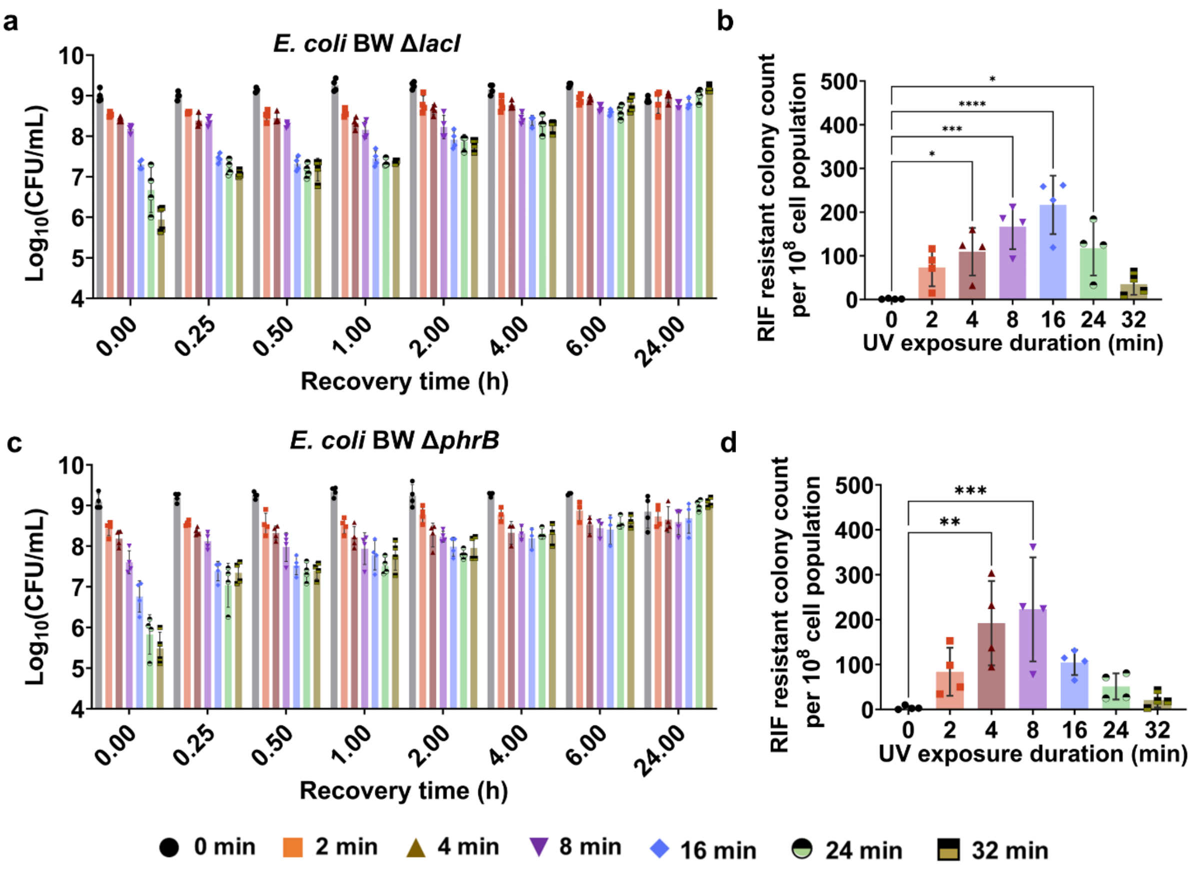

**Fig. S2. Photoreactivation-deficient strain *E. coli* K-12 BW25113 Δ*phrB* exhibited similar trends in survival and mutagenesis compared to the Δ*lacI* control strain.** (a, c) Exponential-phase *E. coli* BW25113 Δ*lacI* and Δ*phrB* cells were exposed to UV-B radiation for 0, 2, 4, 8, 16, 24, and 32 minutes, followed by a 24-hour recovery period. At designated time points, cells were collected and plated to determine CFU levels. (b, d) UV-induced mutagenesis was assessed by quantifying RIF-resistant colonies (per $10^8$ cells) in each knockout strain after recovery at the indicated UV exposure times. Δ*lacI* was used as a reference control. n=4. Statistical analysis was performed using one-way ANOVA with Dunnett's post-test, where $*P < 0.05$, $**P < 0.001$, $***P < 0.01$, $****P < 0.0001$. Data corresponding to each time point represent mean value ± standard deviation.

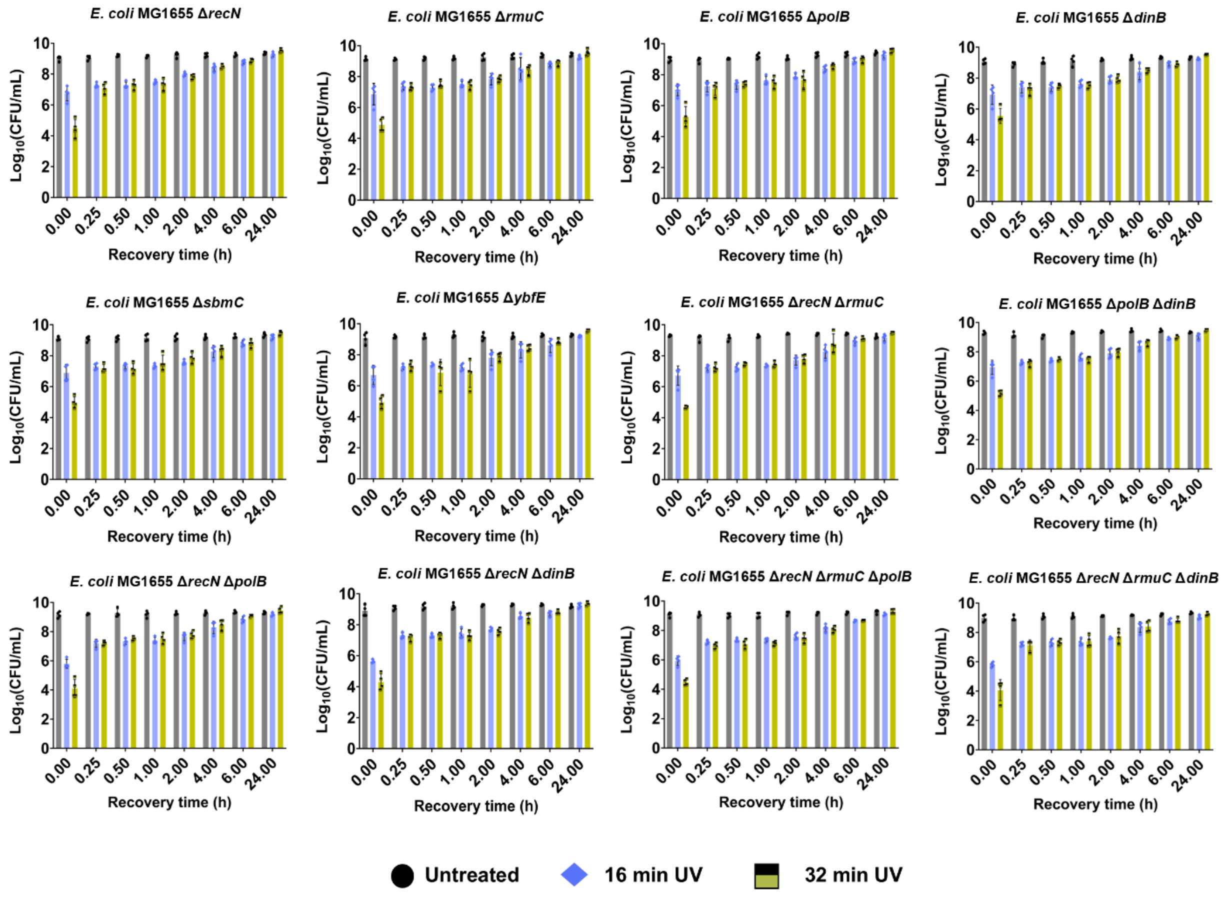

**Fig. S3. Assessment of cell culturability over 24 hours in *E. coli* MG1655 strains lacking UV-upregulated genes identified in the promoter screening (Fig. 2a).** Genes that were upregulated following UV exposure in the MG1655 promoter-reporter library were individually or combinatorially deleted, and UV-induced mutagenesis assays were performed in the resulting knockout strains. Mid-exponential phase cells were exposed to UV-B for 0, 16, or 32 minutes and allowed to recover for 24 hours. At specified recovery time points (0, 0.25, 0.5, 1, 2, 4, 6, and 24 h), samples were collected and plated to assess CFU. n = 4. Data represent mean ± standard deviation for each time point.

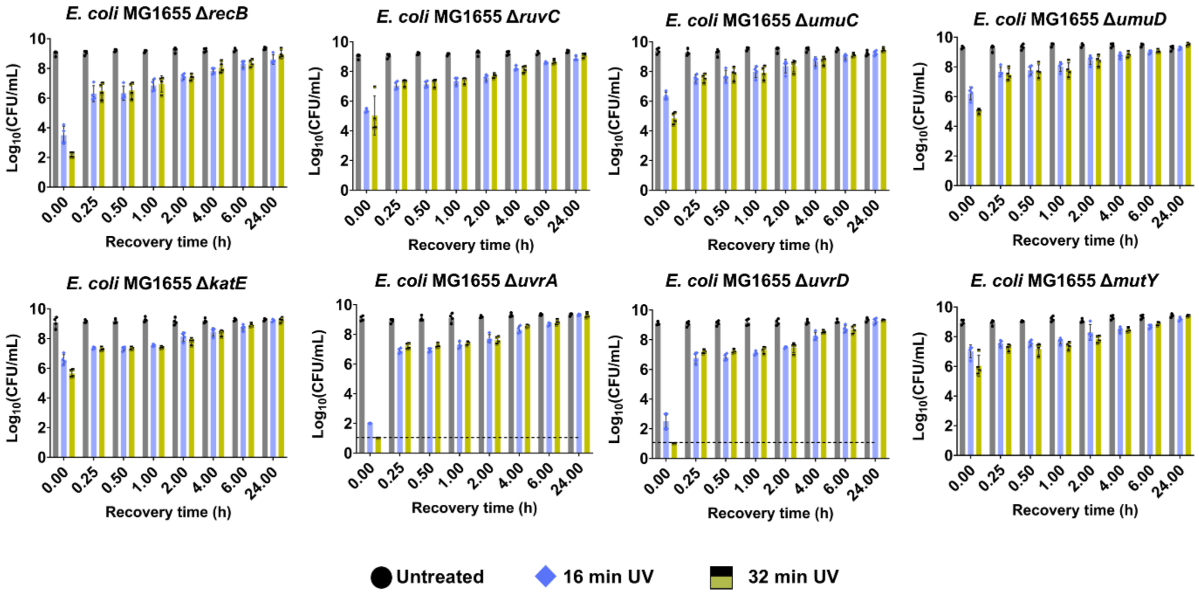

**Fig. S4. Assessment of cell culturability over 24 hours in *E. coli* MG1655 strains lacking selected genes from knockout library screening (Fig. 3a)**. Selected genes identified in the *E. coli* BW25113 single-gene deletion screen were individually deleted in the MG1655 background, and UV-induced mutagenesis assays were performed. Mid-exponential phase cells were exposed to UV-B for 0, 16, or 32 minutes, followed by recovery for 24 hours. At designated time points (0, 0.25, 0.5, 1, 2, 4, 6, and 24 h), samples were collected and plated to assess CFU. n = 4. Data represent the mean ± standard deviation at each time point.

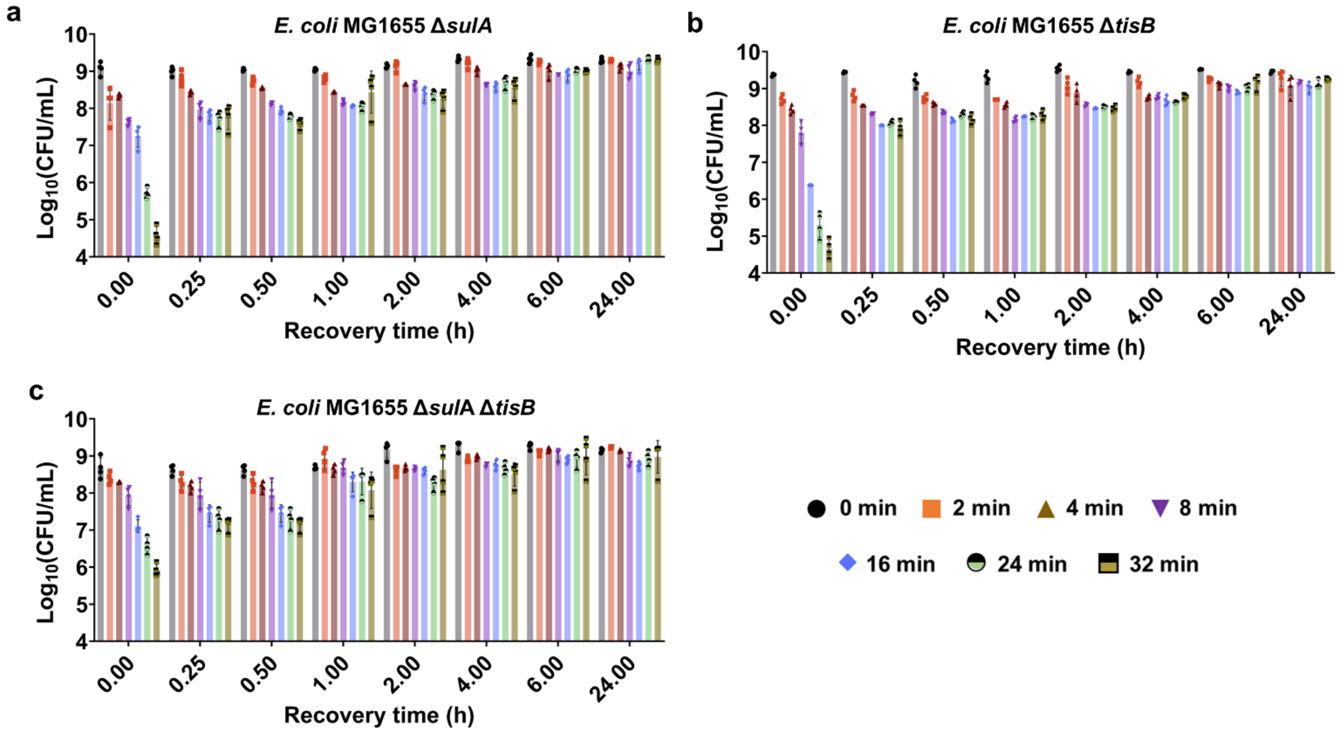

**Fig. S5. Assessment of cell culturability over 24 hours in *E. coli* MG1655 strains lacking SulA and TisB (Fig. 4a-c)**. Mid-exponential-phase *E. coli* MG1655 Δ*sulA*, Δ*tisB*, and Δ*sulA*Δ*tisB* cells were exposed to UV-B light for 0, 2, 4, 8, 16, 24, and 32 minutes, followed by a 24-h recovery period. At designated time points (0, 0.25, 0.5, 1, 2, 4, 6, and 24 h), samples were collected and plated to assess CFU. n = 4. Data represents the mean ± standard deviation at each time point.

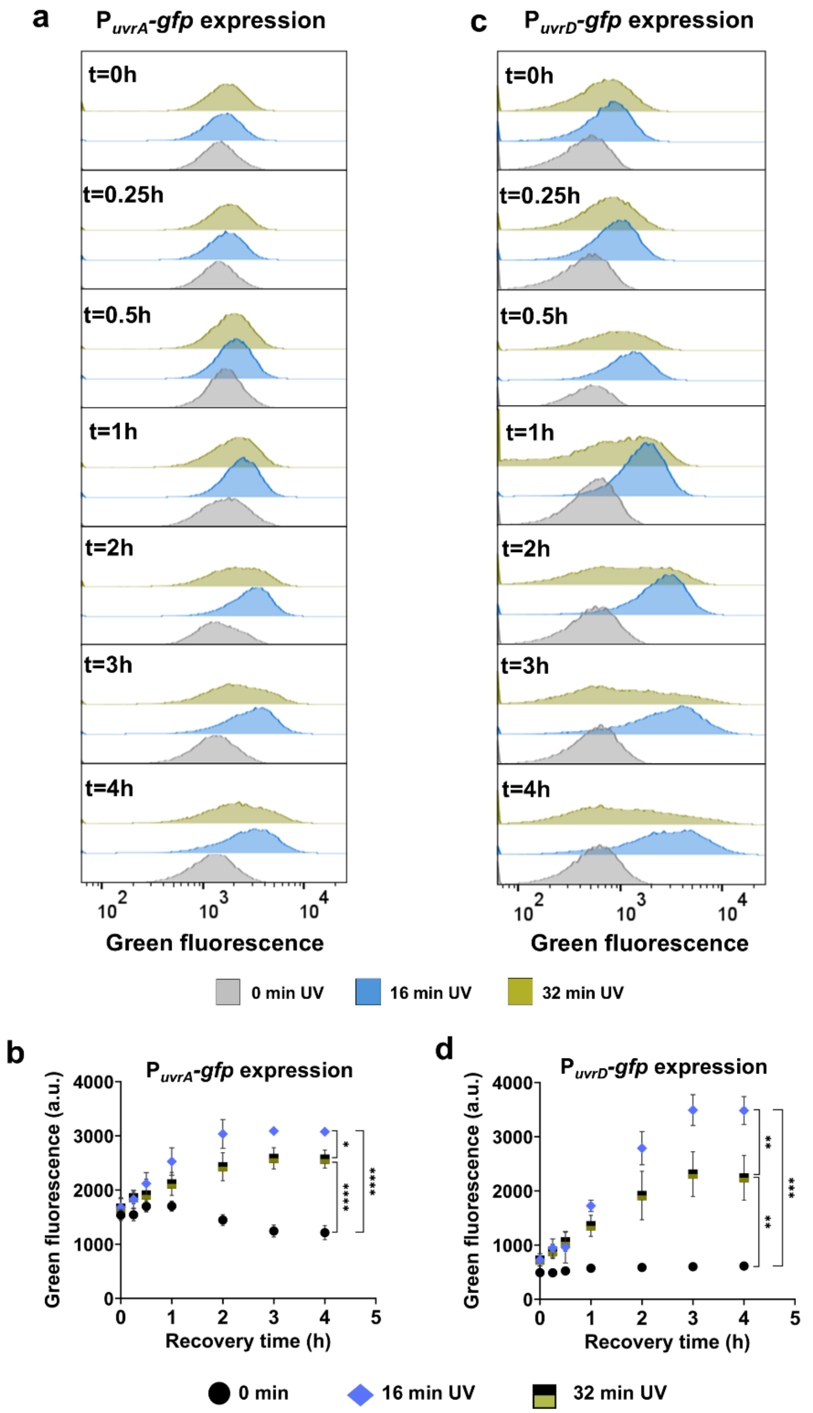

**Fig. S6: Flow cytometry analysis of nucleotide excision repair genes followed by UV-treatment** (a, b) The GFP profiles of UV-treated *E. coli* MG1655 cells harboring *pMSs201* $P_{uvrA}$-*gfp*, and (c, d) the GFP profiles of UV-treated *E. coli* MG1655 cells harboring *pMSs201* $P_{uvrD}$-*gfp* were assessed using flow cytometry after UV exposure (for the indicated time points during recovery). n=4. Statistical analysis was performed using two-way ANOVA, where $*P < 0.05$, $**P < 0.001$, $***P < 0.01$, $****P < 0.0001$. Data corresponding to each time point represent mean value ± standard deviation.

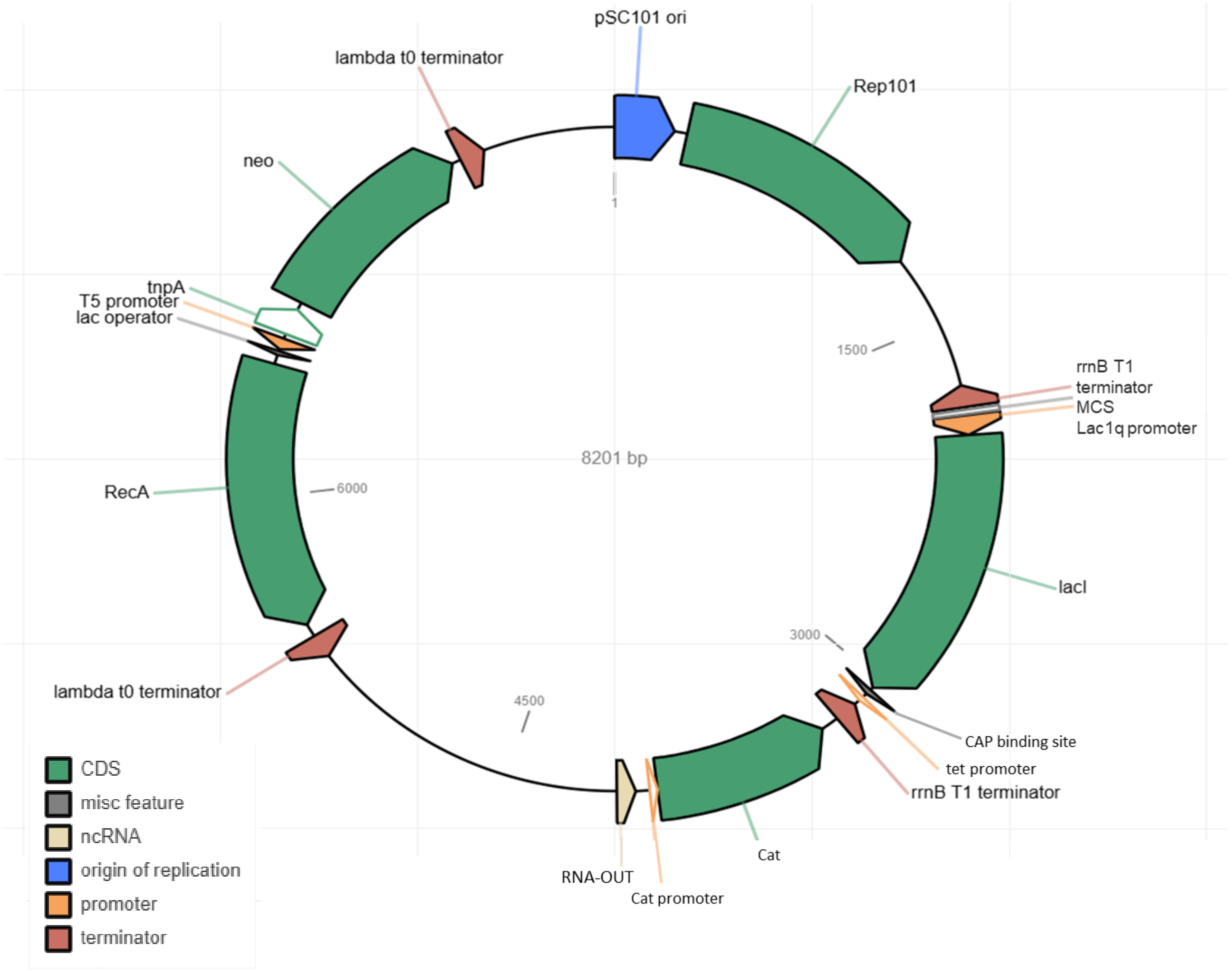

**Fig. S7. Annotated plasmid map of the overexpression plasmid pUA66-*recA*.** An IPTG-inducible *recA* overexpression plasmid was constructed by a commercial cloning service (Synbio Technologies, USA). The *recA* coding sequence from *E. coli* MG1655 was inserted downstream of a T5 promoter in the low-copy plasmid backbone pUA66, which includes a strong mutated *lacI* repressor for tight regulation and a kanamycin resistance marker. The plasmid was verified by sequencing (Plasmidsaurus, USA) and transformed into MG1655 Δ*recA* cells for inducible expression studies.

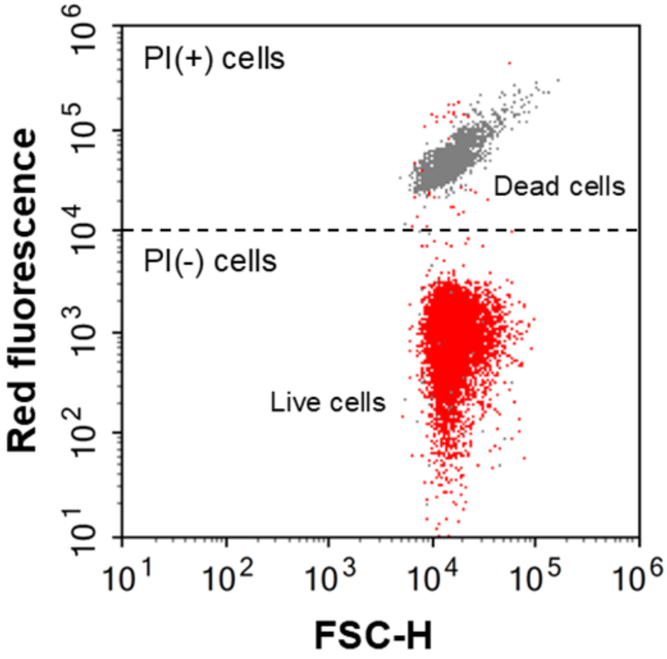

**Fig. S8. PI staining of live and dead cells.** Live cells (Red) and ethanol-treated dead cells (70% v/v, Grey) were stained with PI and analyzed by flow cytometry to assess live and dead cell populations. The figure shows a representative flow cytometry diagram. Similar results were observed across all independent biological replicates, n=4.

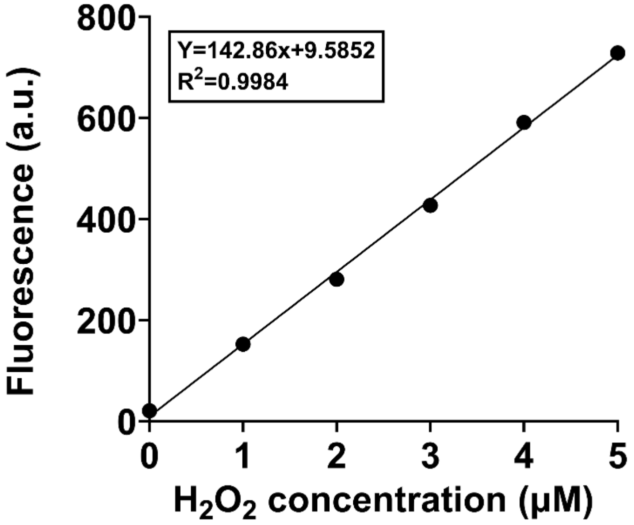

**Fig. S9. Standard curve for $H_2O_2$ measurement using the Amplex Red Hydrogen Peroxide/Peroxidase Assay Kit.** The standard curve was generated using a 20 mM Hydrogen Peroxide ($H_2O_2$) working solution, prepared by dissolving 3.0% (0.88 M) $H_2O_2$ in 1X Reaction Buffer. This solution was then diluted in 1X Reaction Buffer to achieve $H_2O_2$ concentrations ranging from 0 to 10 μM, in 50 μL aliquots. To each microplate well containing the standards, 50 μL of Amplex® Red reagent/HRP working solution was added. After a 30-minute incubation, fluorescence was measured using a microplate reader with excitation in the range of 530–560 nm and emission detection at ~590 nm.

### Supplementary tables

#### Table S1: Promoter and single-deletion knockout libraries used in this study

| Bacterial Strains | Source |
|---|---|
| *E. coli* K-12 MG1655 *pMSs201* $P_{uvrA}$-*gfp* | Promoter library from Horizon Discovery, Lafayette, CO, USA |
| *E. coli* K-12 MG1655 *pMSs201* $P_{dps}$-*gfp* | |
| *E. coli* K-12 MG1655 *pMSs201* $P_{soxS}$-*gfp* | |
| *E. coli* K-12 MG1655 *pMSs201* $P_{recX}$-*gfp* | |
| *E. coli* K-12 MG1655 *pMSs201* $P_{molR}$-*gfp* | |
| *E. coli* K-12 MG1655 *pMSs201* $P_{polA}$-*gfp* | |
| *E. coli* K-12 MG1655 *pMSs201* $P_{dinG}$-*gfp* | |
| *E. coli* K-12 MG1655 *pMSs201* $P_{mutS}$-*gfp* | |
| *E. coli* K-12 MG1655 *pMSs201* $P_{mutH}$-*gfp* | |
| *E. coli* K-12 MG1655 *pMSs201* $P_{mutY}$-*gfp* | |
| *E. coli* K-12 MG1655 *pMSs201* $P_{ybfE}$-*gfp* | |
| *E. coli* K-12 MG1655 *pMSs201* $P_{sbmC}$-*gfp* | |
| *E. coli* K-12 MG1655 *pMSs201* $P_{mutM}$-*gfp* | |
| *E. coli* K-12 MG1655 *pMSs201* $P_{yjiW}$-*gfp* | |
| *E. coli* K-12 MG1655 *pMSs201* $P_{mutT}$-*gfp* | |
| *E. coli* K-12 MG1655 *pMSs201* $P_{dinB}$-*gfp* | |
| *E. coli* K-12 MG1655 *pMSs201* $P_{ftsK}$-*gfp* | |
| *E. coli* K-12 MG1655 *pMSs201* $P_{sodB}$-*gfp* | |
| *E. coli* K-12 MG1655 *pMSs201* $P_{recN}$-*gfp* | |
| *E. coli* K-12 MG1655 *pMSs201* $P_{rmuC}$-*gfp* | |
| *E. coli* K-12 MG1655 *pMSs201* $P_{dinJ}$-*gfp* | |
| *E. coli* K-12 MG1655 *pMSs201* $P_{yebG}$-*gfp* | |
| *E. coli* K-12 MG1655 *pMSs201* $P_{sulA}$-*gfp* | |
| *E. coli* K-12 MG1655 *pMSs201* $P_{soxR}$-*gfp* | |
| *E. coli* K-12 MG1655 *pMSs201* $P_{ahpC}$-*gfp* | |
| *E. coli* K-12 MG1655 *pMSs201* $P_{katE}$-*gfp* | |
| *E. coli* K-12 MG1655 *pMSs201* $P_{sodC}$-*gfp* | |
| *E. coli* K-12 MG1655 *pMSs201* $P_{ahpF}$-*gfp* | |
| *E. coli* K-12 MG1655 *pMSs201* $P_{yhiL}$-*gfp* | |
| *E. coli* K-12 MG1655 *pMSs201* $P_{uvrC}$-*gfp* | |
| *E. coli* K-12 MG1655 *pMSs201* $P_{oxyR}$-*gfp* | |
| *E. coli* K-12 MG1655 *pMSs201* $P_{polB}$-*gfp* | |
| *E. coli* K-12 MG1655 *pMSs201* $P_{uvrD}$-*gfp* | |
| *E. coli* K-12 MG1655 *pMSs201* $P_{sodA}$-*gfp* | |
| *E. coli* K-12 MG1655 *pMSs201* $P_{lexA}$-*gfp* | |

| | |
|---|---|
| *E. coli* K-12 MG1655 *pMSs201* $P_{recA}$-*gfp* | |
| *E. coli* K-12 MG1655 *pMSs201* $P_{lon}$-*gfp* | |
| *E. coli* K-12 MG1655 *pMSs201* $P_{umuD}$-*gfp* | |
| *E. coli* K-12 MG1655 *pMSs201* $P_{ssb}$-*gfp* | |
| *E. coli* K-12 BW25113 Δ*fnr* | |
| *E. coli* K-12 BW25113 Δ*soxR* | |
| *E. coli* K-12 BW25113 Δ*soxS* | |
| *E. coli* K-12 BW25113 Δ*umuD* | |
| *E. coli* K-12 BW25113 Δ*recA* | |
| *E. coli* K-12 BW25113 Δ*ruvB* | |
| *E. coli* K-12 BW25113 Δ*uvrD* | |
| *E. coli* K-12 BW25113 Δ*sodB* | |
| *E. coli* K-12 BW25113 Δ*sodC* | |
| *E. coli* K-12 BW25113 Δ*ahpF* | |
| *E. coli* K-12 BW25113 Δ*katG* | |
| *E. coli* K-12 BW25113 Δ*sodA* | |
| *E. coli* K-12 BW25113 Δ*ygjF* | |
| *E. coli* K-12 BW25113 Δ*polA* | |
| *E. coli* K-12 BW25113 Δ*ruvA* | |
| *E. coli* K-12 BW25113 Δ*ruvC* | |
| *E. coli* K-12 BW25113 Δ*umuC* | |
| *E. coli* K-12 BW25113 Δ*uvrA* | Keio collection, Cat# OEC4987 |
| *E. coli* K-12 BW25113 Δ*alkA* | |
| *E. coli* K-12 BW25113 Δ*ahpC* | |
| *E. coli* K-12 BW25113 Δ*sulA* | |
| *E. coli* K-12 BW25113 Δ*katE* | |
| *E. coli* K-12 BW25113 Δ*uvrB* | |
| *E. coli* K-12 BW25113 Δ*nei* | |
| *E. coli* K-12 BW25113 Δ*dinB* | |
| *E. coli* K-12 BW25113 Δ*recB* | |
| *E. coli* K-12 BW25113 Δ*nth* | |
| *E. coli* K-12 BW25113 Δ*uvrC* | |
| *E. coli* K-12 BW25113 Δ*fur* | |
| *E. coli* K-12 BW25113 Δ*ligB* | |
| *E. coli* K-12 BW25113 Δ*ung* | |
| *E. coli* K-12 BW25113 Δ*tag* | |
| *E. coli* K-12 BW25113 Δ*recN* | |
| *E. coli* K-12 BW25113 Δ*mutY* | |
| *E. coli* K-12 BW25113 Δ*ycaQ* | |

| |
|---|
| *E. coli* K-12 BW25113 Δ*mutM* |
| *E. coli* K-12 BW25113 Δ*recC* |
| *E. coli* K-12 BW25113 Δ*polB* |
| *E. coli* K-12 BW25113 Δ*recD* |
| *E. coli* K-12 BW25113 Δ*oxyR* |
| *E. coli* K-12 BW25113 Δ*mutS* |
| *E. coli* K-12 BW25113 Δ*mutH* |
| *E. coli* K-12 BW25113 Δ*mutL* |
| *E. coli* K-12 BW25113 Δ*exoI* |
| *E. coli* K-12 BW25113 Δ*phrB* |
| *E. coli* K-12 BW25113 Δ*lacI* |

**Table S2: Bacterial plasmids and strains used in this study**

| **Bacterial Strains** | **Source** |
|---|---|
| *E. coli* K-12 MG1655 pUA66 P*recA-gfp* | Gift from Dr. Mark P. Brynildsen |
| *E. coli* K-12 MG1655 pUA66-*gfp* | Gift from Dr. Mark P. Brynildsen |
| *E. coli* K-12 MG1655 Wild type | Gift from Dr. Mark P. Brynildsen |
| *E. coli* K-12 MG1655 Δ*sulA* | Previous study |
| *E. coli* K-12 MG1655 Δ*tisB* | Previous study |
| *E. coli* K-12 MG1655 Δ*recA* | Previous study |
| *E. coli* K-12 MG1655 Δ*sulA*Δ*tisB* | This study |
| *E. coli* K-12 MG1655 Δ*recN* | This study |
| *E. coli* K-12 MG1655 Δ*rmuC* | This study |
| *E. coli* K-12 MG1655 Δ*polB* | This study |
| *E. coli* K-12 MG1655 Δ*dinB* | This study |
| *E. coli* K-12 MG1655 Δ*sbmC* | This study |
| *E. coli* K-12 MG1655 Δ*ybfE* | This study |
| *E. coli* K-12 MG1655 Δ*polB*Δ*dinB* | This study |
| *E. coli* K-12 MG1655 Δ*recN*Δ*rmuC* | This study |
| *E. coli* K-12 MG1655 Δ*recN*Δ*polB* | This study |
| *E. coli* K-12 MG1655 Δ*recN*Δ*dinB* | This study |
| *E. coli* K-12 MG1655 Δ*recN*Δ*rmuC*Δ*polB* | This study |
| *E. coli* K-12 MG1655 Δ*recN*Δ*rmuC*Δ*dinB* | This study |
| *E. coli* K-12 MG1655 Δ*recB* | This study |
| *E. coli* K-12 MG1655 Δ*ruvC* | This study |
| *E. coli* K-12 MG1655 Δ*umuC* | This study |
| *E. coli* K-12 MG1655 Δ*umuD* | This study |

| | |
|---|---|
| *E. coli* K-12 MG1655 Δ*katE* | This study |
| *E. coli* K-12 MG1655 Δ*uvrA* | This study |
| *E. coli* K-12 MG1655 Δ*uvrD* | Previous study |
| *E. coli* K-12 MG1655 Δ*mutY* | This study |
| *E. coli* K-12 MG1655 Δ*recA*+pUA66-*recA* | This study |
| *E. coli* K-12 MG1655 Δ*recA*+ pUA66-E.V. | This study |

**Table S3: Oligonucleotides used to generate single and multi-deletion mutants**

| **Oligonucleotides to generate gene deletions** | | | |
|---|---|---|---|
| **Mutation** | **Forward Primer (5’ to 3’):** | **Reverse Primer (5’ to 3’)** | **Source** |
| Δ*recN*::KAN$^R$ | ACACAATAACAGTAATGG<br>TTTTTCATACAGGAAAAC<br>GACTGTGTAGGCTGGAGC<br>TGCTTC | CGTTTTGCTGTTTACTC<br>TGACCGTGAAGCAGGA<br>AAAAAGTTTAACGGCT<br>GACATGGGAAT | Integrated DNA Technologies, Inc. |
| Δ*rmuC*::KAN$^R$ | CAGGAAATGCCTTTCCA<br>ACTGGACGTTTGTACAG<br>CACAATTCTATTTTGTG<br>CGGGTAAGTTGTTGCGT<br>CAGGAGGCGTTGTGTA<br>GGCTGGAGCTGCTTC | TCAAAAAATTGTTCC<br>AGAAGTGTAACAGAT<br>TGGGCGTCGATGCCC<br>TAGATTTCTACCCGG<br>CTTAACTACTCCCAA<br>TGGGTTAACGGCTGA<br>CATGGGAAT | Integrated DNA Technologies, Inc. |
| Δ*polB*::KAN$^R$ | CAGGCTATACTCAAGCC<br>TGGTTTTTTGATGGATT<br>TTCAGCGTGTAGGCTGG<br>AGCTGCTTC | TCACGCATCAAAATG<br>GTATCTGGCGAACTC<br>TTTTTTTTGCTTAACG<br>GCTGACATGGGAAT | Integrated DNA Technologies, Inc. |
| Δ*dinB*::KAN$^R$ | ACGCGTTAAATGCTGA<br>ATCTTTACGCATTTCTC<br>AAACCCTGAAATCACT<br>GTATACTTTA<br>CCAGTGTTGAGAGGTG<br>AGCA<br>GTGTAGGCTGGAGCTG<br>CTTC | CCGATTTTTCAGCGA<br>GAATTCGATGCATAC<br>AGTGATACCCTCATA<br>ATAATGCACACCAGA<br>ATATACATAATAGTA<br>TACATTAACGGCTGA<br>CATGGGAAT | Integrated DNA Technologies, Inc. |
| Δ*sbmC*::KAN$^R$ | CAACTATACTGTATATA<br>AAAACAGTATCAATGG<br>AGGCGTCGTGTAGGCT<br>GGAGCTGCTTC | AAAGAGTGGTCATCG<br>CGTTAACACACCGCC<br>CTGAGATGAATTAAC<br>GGCTGACATGGGAA | Integrated DNA Technologies, Inc. |
| Δ*ybfE*::KAN$^R$ | AGAGGCCTGGCTGATT<br>GTTTCCCCCGAAGTCAC<br>CAAGATCGTGTAGGCT<br>GGAGCTGCTTC | ACGAGCAGGACTGCA<br>CACTGTGCTACATGA<br>AAGTGGAAATTTAAC<br>GGCTGACATGGGAAT | Integrated DNA Technologies, Inc. |

| | | | |
|---|---|---|---|
| Δ*recB*::KAN$^R$ | AGCGCGTTGCAGCAAACAATGCCCCTGATGAGTGAAAAGAGTGTAGGCTGGAGCTGCTTC | GCGGGCGTAGCTGTTTGTGCTCCACAGCTTCCAGTAATTGCTTTTGCAATTTCATTAACGGCTGACATGGGAAT | Integrated DNA Technologies, Inc. |
| Δ*ruvC*::KAN$^R$ | CTCTGATGAGGCCTGCTAAACAGCAAAACGGAGACGCGTGGTGTAGGCTGGAGCTGCTTC | GTGGAAACGCCTCAGCCGGAACTGACCGAGGCGGTATAACTTAACGGCTGACATGGGAAT | Integrated DNA Technologies, Inc. |
| Δ*umuC*::KAN$^R$ | TCTTTGGTGTGGTGATCCACGTCGTTAAGGCGATGCGCTGGTGTAGGCTGGAGCTGCTTC | TCGGCGCTCCTGCGGGAGCGCTTTTTCCTGCCGCTATATTTAACGGCTGACATGGGAA | Integrated DNA Technologies, Inc. |
| Δ*umuD*::KAN$^R$ | GAACAGACTACTGTATATAAAAACAGTATAACTTCAGGCAGATTATTGTGTAGGCTGGAGCTGCTTC | CAGCTGGCATAAAACGCGTTTACATCACAGAGGGCAAACATTTAACGGCTGACATGGGAAT | Integrated DNA Technologies, Inc. |
| Δ*katE*::KAN$^R$ | ACAGCGGCCCTTTCAGTAATAAATTAAGGAGACGAGTTCAGTGTAGGCTGGAGCTGCTTC | ATGTAAATCATTTGAGGCGGCGCAATTGCGCCGCCTCCCATTAACGGCTGACATGGGAA | Integrated DNA Technologies, Inc. |
| Δ*uvrA*::KAN$^R$ | ATGCCACCGGGCAAAAAAGCGTTTAATCCGGGAAAGGTGAGTGTAGGCTGGAGCTGCTTC | CTCTGAAAGGAAAAGGCCGCTCAGAAAGCGGCCTTAACGATTAACGGCTGACATGGGAA | Integrated DNA Technologies, Inc. |
| Δ*mutY*::KAN$^R$ | TGCTGCAATCTTGCCCCCAACAACAGTGAATTCGGTGACCGTGTAGGCTGGAGCTGCTTC | TCGTTCTGCTCATAAATCATCCTCTTTATCGACTCACGCGTTAACGGCTGACATGGGAAT | Integrated DNA Technologies, Inc. |
| Δ*phrB*::KAN$^R$ (Trial 1) | CTTTGGCCGCGTGTGATTAACTTGCGCCATTCAGGAGTTTTGTGTAGGCTGGAGCTGCTTC | CAGACGCGTCAGGCAATCGAGCCCAGATGCCGGATGCGGCTTAACGGCTGACATGGGAAT | Integrated DNA Technologies, Inc. |
| Δ*phrB*::KAN$^R$ (Trial 2) | AGGGCAGCGTTATTTCGAACTTTGGCCGCGTGTGATTAACTTGCGCCATTCAGGAGTTTTGTGTAGGCTGGAGCTGCTTC | CTGTTCGGTCGCTAATCCATTCGGCGCTCCTGCGGGAGCGCTTTTTTCCTGCCGCTATATTTAACGGCTGACATGGGAAT | Integrated DNA Technologies, Inc. |

**Oligonucleotides to verify gene deletions**

| Mutation | External Forward Primer (5’ to 3’) | External Reverse Primer (5’ to 3’) | Internal Forward Primer (5’ to 3’) | Internal Reverse Primer (5’ to 3’) | Source |
|---|---|---|---|---|---|

| | | | | | |
|---|---|---|---|---|---|
| Δ*recN*::KAN$^R$ | GATTCGTC GCTGTGAT TACCATC | GCTCTTCGT CCAGATCAT CCT | GCACAACT GACCATCA GCAA | CAGTTGC TGCTGTT CTTCCAG TAG | Integrated DNA Technologies, Inc. |
| Δ*rmuC*::KAN$^R$ | AAAGCCA TGCGGTG AAAATC | GCTCTTC GTCCAGA TCATCCT | GTTATTGC GTTGGTGG GTGT | GGGTTCA ACGGGA ATAAACA | Integrated DNA Technologies, Inc. |
| Δ*polB*::KAN$^R$ | GTCAGTTA GCGCCGC AGTTA | GCTCTTC GTCCAGA TCATCCT | GCGCAGGC AGGTTTTA TCTT | GGTTTAT CTTCGGC GAAACG | Integrated DNA Technologies, Inc. |
| Δ*dinB*::KAN$^R$ | GACCAAA AGTGCGTC CGATA | GCTCTTC GTCCAGA TCATCCT | GTGGATAT GGACTGCT TTTTCG | GGCGTTC ATCCCAG GTTTTA | Integrated DNA Technologies, Inc. |
| Δ*sbmC*::KAN$^R$ | GGTTACCC TGGAATCT GTCG | GCTCTTC GTCCAGA TCATCCT | GAAGAGA AACGTACC GTTGCAG | GCTGCAC CGCAACA TACATT | Integrated DNA Technologies, Inc. |
| Δ*ybfE*::KAN$^R$ | TACGCGCT ATCCGTCG CTAT | GCTCTTC GTCCAGA TCATCCT | AACAAACG GACCGTAC GACA | GCCAGTT GCTGCAT TAACATC T | Integrated DNA Technologies, Inc. |
| Δ*recB*::KAN$^R$ | TTCTTCCA TCAGGCG GTGGT | GCTCTTC GTCCAGA TCATCCT | ATCCACTG TACGAACG CCTG | AAATTCG GTACTGC TGGGGG | Integrated DNA Technologies, Inc. |
| Δ*ruvC*::KAN$^R$ | GAAACCG CACCGAA ACTGAT | GCTCTTC GTCCAGA TCATCCT | GGCTATTA TTCTCGGC ATTGATCC | AACGTGG CAGTGGG TGATAG | Integrated DNA Technologies, Inc. |
| Δ*umuC*::KAN$^R$ | CTGTTGAC GGCGAGT TTACG | GCTCTTC GTCCAGA TCATCCT | AGCTGTGA GACGGTGT TTCG | GATCGCG TAGCAGC GTTAAT | Integrated DNA Technologies, Inc. |
| Δ*umuD*::KAN$^R$ | TTTTCATT TCTGCTGG ATGAGC | GCTCTTC GTCCAGA TCATCCT | TCTTGTTC AGTGTGGC TTTCC | TGGATCA CCACACC AAAGAC A | Integrated DNA Technologies, Inc. |
| Δ*katE*::KAN$^R$ | GTTTAGCC GATTTAGC CCCTG | GCTCTTC GTCCAGA TCATCCT | ATCAGCCG CTCACGGT TATT | TATGGTA AGGGCA GGTCGGA | Integrated DNA Technologies, Inc. |

| | | | | | |
|---|---|---|---|---|---|
| Δ*uvrA*::$KAN^R$ | F_Ex_sodA: CACAACA CTCCGGGT AATGC | GCTCTTC GTCCAGA TCATCCT | TGAATCCC TTTCCGCC TACG | ACGGATG ACGACGA ATGGAG | Integrated DNA Technologies, Inc. |
| Δ*mutY*::$KAN^R$ | CAAATTCC GGTGAAA TGACG | GCTCTTC GTCCAGA TCATCCT | GCCCAGGT TCTGGACT GGTA | CGACACG GGAAGCC ACATAG | Integrated DNA Technologies, Inc. |
| Δ*phrB*::$KAN^R$ (Trial 1) | TCCTGACG CCTGGCTT TCAG | GCTCTTC GTCCAGA TCATCCT | ATATCGCT ACACCACG CCAG | AACAAGC GATGCAA GCACTG | Integrated DNA Technologies, Inc. |
| Δ*phrB*::$KAN^R$ (Trial 2) | TTCCACAG AACGCCA GCAGC | GCTCTTC GTCCAGA TCATCCT | CCAGTGGG CGACGCAT AACA | AGCGGCA TCAACAA TCGGGT | Integrated DNA Technologies, Inc. |
